# Persistent interferon signaling that causes sensory neuron plasticity and pain in arthritis

**DOI:** 10.1101/2025.01.18.633447

**Authors:** Jie Su, Ming-Dong Zhang, Jussi Kupari, Dongoh Kwak, Laurence Picton, Bingze Xu, Yizhou Hu, Alejandro Gonzalez Alvarez, Dmitry Usoskin, Zhongwei Xu, Abdeljabbar El Manira, Rikard Holmdahl, Patrik Ernfors

## Abstract

While the inflammatory processes in rheumatoid arthritis have been described, mechanisms driving pain are poorly defined. Here, we used a multitude of approaches to uncover the neural basis, mediators, intracellular signaling pathway and the mechanism of inflammatory pain. In cartilage autoantibody-induced arthritis mice, an early immune-activation and a cytokine storm were mainly driven by vascular cells and monocyte/macrophages in the dorsal root ganglion. However, persistently elevated interferons and receptor-activation of the MNK1/2-eIF4E signaling pathway at all disease phases caused sensory-motor dysfunction and pain by inducing hyperexcitability and sensitization of *Gfra3*+ sensory neurons. Like mice, human sensory neurons expressed interferon receptors and interferons were elevated only in individuals with painful rheumatoid arthritis. Signaling pathway inhibition *in vivo* reversed pain and restored limb function. The finding that joint pain before and during arthritis is caused by a defined cytokine and signaling pathway holds promise for targeted therapies for pain relief in arthritis.

## Introduction

Emerging data has shown that in many inflammatory conditions, pain develops before onset of inflammation and persists even after inflammation has resolved indicating that it is not merely an accompanying symptom of inflammation^1- 3^. Joint pain, arthralgia, is typically appearing before the onset of rheumatoid arthritis (RA) and remains irrespective of the severity of arthritis. An efficient treatment of arthritis is commonly not curing pain symptoms, i.e. “remaining pain” affects as many as 25% of patients and may result in considerable suffering^1-3^.

Pain associated with inflammation has been ascribed to an increase in excitability of sensory neurons residing in dorsal root ganglia^4, 5^. Activity in primary sensory neurons is essential, because local anaesthesia relieves not only acute but also chronic pain^6, 7^. RA is associated with a strong autoantibody response and involves production of several cytokines of different classes, and some of these contribute to the inflammatory disease, such as tumor necrosis factor (TNF), IL-1β and IL-6. Both cartilage binding autoantibodies and several inflammatory cytokines cause dorsal root ganglion (DRG) neuron hyperexcitability and sensitizes to painful stimuli when injected into the experimental animal^8-13^. Various alternative mediators have also been suggested to induce hyperexcitability and pain including for example, ligands for toll-like receptors, inflammatory lipid mediators, neuropeptides and growth factors^14-22^. In mouse, many different types of DRG sensory neuron types generating pain have been identified by single-cell RNA-sequencing (scRNA-seq) and the human evolutionary cell type homologs are known^23-27^. Most of these express neuropeptides consistent with peptidergic nerve fibers localized to joint tissues^28, 29^, although little is known which sensory neuron types are functionally involved and how they are molecularly perturbed in arthritis. Thus, the importance of the various inflammatory mediators, the identity of the pain causing sensory neuron population and the intracellular mechanisms causing hyperexcitability and pain in arthritis is not fully understood.

Type I interferons (IFN1) orchestrate the body’s defense predominantly against viruses. While serum levels of IFN1 are increased at early phases of RA, they are not considered essential for development of arthritis^4, 30^. We reasoned that by identifying the neuron type that causes pain, and characterizing the intercellular communication between non-neuronal, immune cells and neurons, we could identify the cellular and molecular mechanisms responsible for sensory dysfunction in arthritis. Our work reveals that hyperexcitability in *Gfra3*^+^ sensory neurons is caused by an alternative type I interferon signaling pathway resulting in joint pain and impairment of dexterity and overall limb function.

## Results

### A sensitization of a variety of sensory neuron types in autoantibody-induced arthritis and post-arthritis mice

We used a cocktail of monoclonal pathogenic antibodies targeting cartilage, similar to antibodies present in RA to trigger onset of arthritis in mice (the cartilage autoantibody-induced arthritis model)^31^. Macroscopic arthritis (swelling and redness) developed between day 6 and 23 after injection of the autoantibodies directed towards native and citrullinated cartilage proteins, enhanced by LPS injection on day 5 (Fig. 1a,b). The mice developed allodynia within 4 hours with reduced withdrawal threshold to von Frey filaments before any inflammation (4h, d1, d3), during inflammation (d9, d12, d17, d23) as well as long after inflammation had resolved (d30, d40, d46, d63) (Fig. 1c and Extended Data Fig. 1a). Thus, this indicated that the arthritis model captured features of arthralgia before onset of arthritis, as well as post-arthritis remaining pain. To examine this further, we used the inverted screen test as a measure of overall sensory-motor limb function^32^. Animals performed less well in the inverted screen at all three disease phases after injection of the antibodies (Fig. 1d). A firm handshake with arthritis can be painful due to tenderness in the joints. We devised a paw squeeze test which revealed a marked increase in pain-like behavior in autoantibody injected animals, indicating arthralgia prior, during and after inflammation had resolved (i.e. remaining post-arthritis pain) (Fig. 1e). Stiffness and loss of fine motor skills were assayed by alterations in the dexterous use of the forepaws assessed by the sunflower seed handling behavioral assay in which the animals are challenged to reach and successfully peel the shell to expose the seed kernel^33^. Control mice grasped and held the sunflower seed between the forepaws and systematically deshelled it by rotating the seed while clamping the upper and lower incisors into the shell surface edge, resulting in the exposure of an intact seed kernel (Fig. 1f and Supplementary Video 1). Mice with arthritogenic autoantibodies, before, during or after the development of arthritis, were unable to grasp and/or maintain a firm grip on the seed, largely failed to rotate the seed for deshelling and instead adapted to several alternative strategies that resulted in partial peeling and a failure to expose an intact seed kernel (Fig. 1f and Supplementary Video 2). The mice also showed increased nocifensive behavior to cutaneous pricking (mechanical hyperalgesia) and cold hyperalgesia before, during and after the period of joint inflammation (Fig. 1g,h). Sensitization emerged as short as 4h after autoantibody injection (Extended Data Fig. 1a). Thus, injection of the autoantibodies leads to a general reduction of limb function, arthralgia, loss in integrity of dexterous use of paws and cutaneous mechanical and cold hyperalgesia before any observed inflammation, during inflammation and persisted after inflammation had resolved, similar to clinical observations in RA^1-3, 34-36^.

**Fig. 1.**
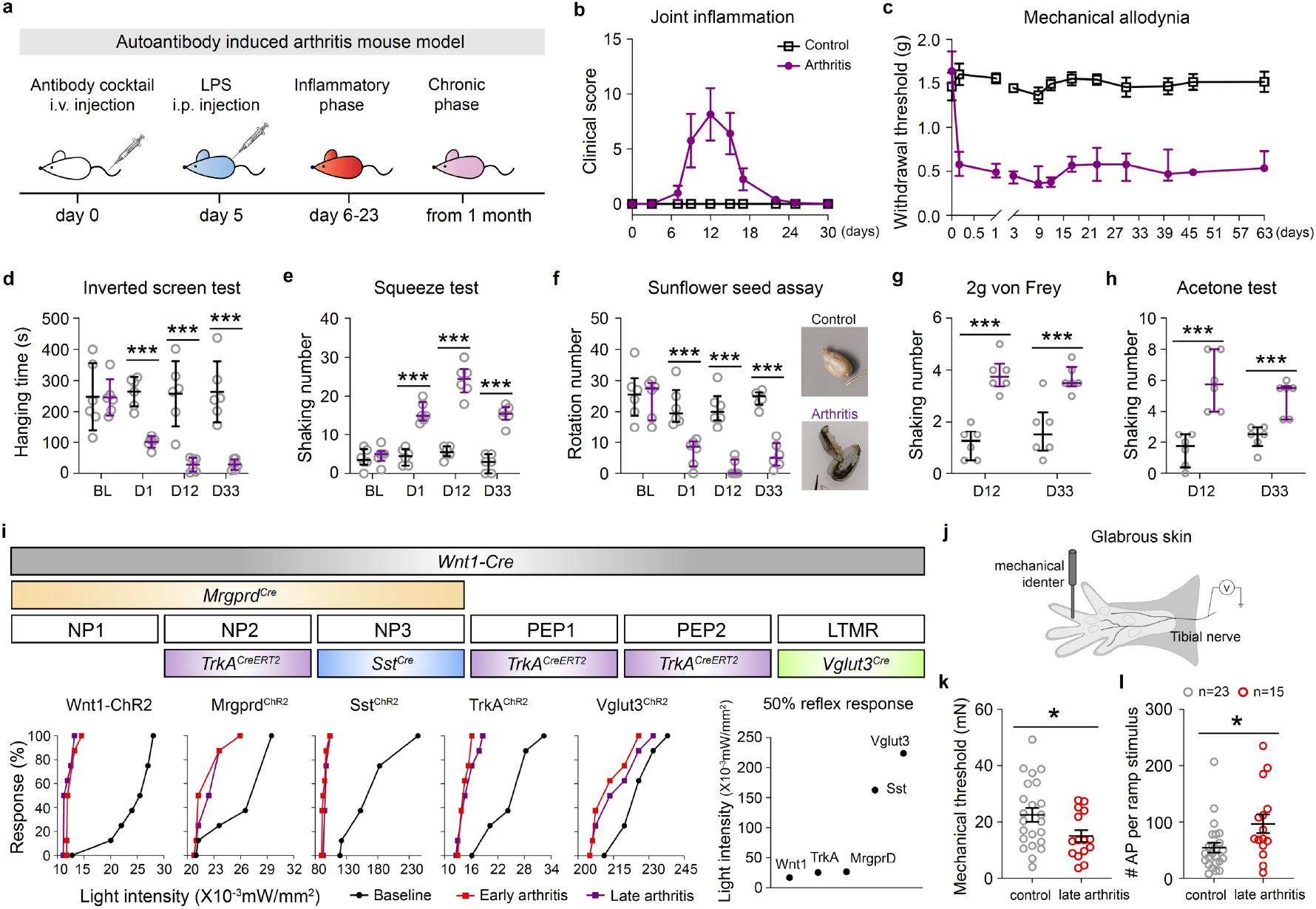
Characterization of pain-like behaviors in autoantibody-induced arthritis model. (**a**) Cartilage autoantibody-induced arthritis mouse model. (**b**) Time course of transient joint inflammation (clinical score) from day 6 to day 23 after autoantibody injection in C57BL/6N mice (n = 6 per group in **b**-**h**). LPS was injected on day 5 to enhance arthritis development. (**c**) Time course of mechanical allodynia in arthritis mice starting as early as 4 hours and lasting until day 63 after autoantibody injection. (**d**) Hanging time of inverted screen test in different timepoints after antibody injection (D1: arthralgia; D12: arthritis with peak inflammation; D33: arthritis after inflammation remission) in control and arthritis mice. BL: baseline control. \*\*\**p* < 0.001. (**e**) Shaking numbers of clip squeeze test after antibody injection, \*\*\**p* < 0.001. (**f**) Rotation numbers of sunflower seed assay and seed images after peeling the shell to expose the seed kernel, \*\*\**p* < 0.001. (**g**) Shaking numbers of 2g von Frey test, \*\*\**p* < 0.001. (**h**) Shaking numbers in acetone test, \*\*\**p* < 0.001. (**i**) Top: Cre and CreERT2 mouse strains used in the study to target specific DRG neuron populations. Bottom: Reflex responses percentage to blue light stimulation in the mouse strains (crossed with R26-ChR2) under different stages after arthritis (Early arthritis: early phase of autoantibody-induced arthritis during inflammation, day 9-21; Late arthritis: late phase of arthritis after remission of inflammation, after day 30); light intensity for inducing reflex response in 50% of mice from different strains (n = 8/strain). (**j**) Skin-nerve recording paradigm. (**k**) Mechanical thresholds of mice injected with saline (“control”) or autoantibody (around 3 months after antibody injection, “late arthritis”). Dots represent values for individual fibers, lines represent the mean and SD. (**l**) Number of mechanically induced action potentials during the force ramp application, \**p* < 0.05. One-way ANOVA followed by Dunnett multiple comparisons test for (**d**), Kruskal-Wallis test followed by Dunn’s multiple comparisons test for (**e**-**h**), unpaired t test for (**j, k**).

We next asked whether autoantibody-induced arthritis leads to a sensitization of sensory nerves. To identify the neuronal types which become sensitized, we used mouse driver lines to target the molecularly different sensory neuron subtypes identified by scRNA-seq analysis^24^. The following Cre mouse lines were used: *Wnt1*-*Cre* with expression in all sensory neurons^37^; *Mrgprd*^*Cre*^ mice with expression in pruriceptors (NP1, NP2 and NP3 neurons)^38^; *TrkA*^*CreERT2*^ mice with expression in C-nociceptors (PEP1), Aδ-heat nociceptors (PEP2), Aδ-high threshold mechanoreceptor (PEP3) and *Mrgpra3*^+^ pruriceptors (NP2 neurons)^39^; *Sst*^*Cre*^ mice targeting NP3 pruriceptors^40^; and *Vglut3*^*Cre*^ mice targeting C-low threshold mechanoreceptors^41^ (Fig. 1i). We confirmed patterns of cell type-specific recombination consistent with previous characterizations by crossing the driver lines with a *ROSA26*^*Tomato*^ reporter strain (Extended Data Fig. 1b) and thereafter crossed them with Ai32 mice carrying a conditional ChR2 allele, hereafter referred to as Wnt1-ChR2, Mrgprd^ChR2^, TrkA^ChR2^, Sst^ChR2^ and Vglut3^ChR2^ mice. To assess sensitization, increasing intensity of light was applied to the paw of the ChR2 expressing mice and the light-induced withdrawal threshold was recorded. In control mice, a striking difference was observed in the threshold between different sensory neuron types, with Wnt1-ChR2, TrkA^ChR2^ and Mrgprd^ChR2^ mice being most sensitive (13-21×10^−3^ mW/mm^2^) while Sst^ChR2^ and Vglut3^ChR2^ mice required more than 100×10^−3^ mW/mm^2^. Following autoantibody-induced arthritis, all neuron types were sensitized, and the sensitization persisted both in the arthritis and post-arthritis phases. Nevertheless, sensitization varied, with Wnt1-ChR2 and TrkA^ChR2^ mice displaying the greatest shift in light intensity leading to a behavioral response (Fig. 1i). These results indicate that the exposure to autoantibodies leads to a general sensitization and sensory dysfunction indiscriminate of sensory neuron subtypes which could explain the multiple sensory-motor symptoms of RA.

TrkA expressing neurons include the mechanosensitive C-fiber nociceptive neurons^42^. To directly assess action potential threshold and frequency, we performed single nerve recordings of mechanosensitive C-fibers in mice using the skin-nerve preparation (Fig. 1j). Individual C-fibers and the receptive field were identified and a force ramp from 0-100 mN (10 sec) was applied to find the mechanical threshold and to determine the firing frequency during the ramp in control mice and autoantibodies treated mice (Extended Data Fig. 1c). The mechanical threshold was lower in arthritis mice and the average firing frequency (action potentials, APs, per second) and total APs per ramp stimulus was markedly increased, starting already at relatively low forces (Fig. 1k,l and Extended Data Fig. 1c,d). Thereafter 10 sec force steps were applied with the forces: 10, 20, 40, 50, 75, 150, 200 mN (Extended Data Fig. 1e). Consistently, measuring average mechanically induced APs during the force step application revealed increased firing starting already at 40 mN force in arthritis mice (Extended Data Fig. 1f). Thus, we conclude a sensitization involving axon hyperexcitability affecting both activation thresholds and supra-threshold responses.

### Pain in arthritis is caused by a molecularly defined population of sensory neurons

The above results revealed that many kinds of sensory neurons become sensitized, however, a select subset of neuron types is likely to cause allodynia and hyperalgesia in arthritis. To identify the neurons, we combined light activation which by itself did not lead to any behavior (subthreshold) in Wnt1-ChR2, Mrgprd^ChR2^, TrkA^ChR2^, Sst^ChR2^ and Vglut3^ChR2^ mice with von Frey threshold reflex measurements, nocifensive behavior in response to 2g von Frey pricking and to cold (acetone)^43^. We measured basal response prior to arthritis, early (arthralgia with inflammation) and late (post-inflammation) phases of induced arthritis with and without subthreshold light. Injection of autoantibodies led to allodynia and hyperalgesia in all mouse strains, but subthreshold light potentiated allodynia and nocifensive behavior to pricking and cold only in the TrkA^ChR2^ mice and the positive control Wnt1-ChR2 mice which target all sensory neuron types (Fig. 2a). To confirm the critical role of TrkA expressing sensory neurons we crossed the TrkA^CreERT2^ mice to Ai40 mice carrying a conditional ArchT allele (TrkA^ArchT^ mice) to optogenetically silence TrkA^CreERT2^ expressing neurons. Silencing resulted in a complete reversal of allodynia, pricking and cold hyperalgesia both during and after resolution of inflammation (Fig. 2b and Extended Data Fig. 2a). Furthermore, joint pain and deficits in overall limb function and sensory-motor functions were reversed by optogenetic inhibition of TrkA neurons (Fig. 2b). Several different molecularly defined sensory neuron types express TrkA (PEP1, PEP2, PEP3 and NP2). We next set out to establish whether the C-fiber nociceptor neuron type PEP1 among the TrkA expressing sensory neuron types is responsible for pain in arthritis. *Gfra3* is expressed exclusively in PEP1 sensory neurons^24^. We therefore generated *Gfra3*^*CreERT2*^ mice carrying a conditional reporter, ChR2 or ArchT allele (Gfra3^TOM^, Gfra3^ChR2^ and Gfra3^ArchT^ mice) and injected autoantibodies in the latter two strains of mice. Gfra3^TOM^ mice confirmed recombination in the DRG (Extended Data Fig. 2b) and C-mechanonociceptors in Gfra3^ChR2^ mice were sensitized, as shown by reduced light-induced withdrawal thresholds which persisted both during inflammation and after resolution of inflammation (Extended Data Fig. 2c). In Gfra3^ChR2^ mice, subthreshold light markedly potentiated allodynia, pricking and cold pain behavior (Fig. 3a) and silencing the Gfra3^+^ neurons in Gfra3^ArchT^ mice with arthritis reversed allodynia, pricking and cold pain behavior both during inflammation and after resolution (Fig. 3b). In control animals, inhibition of Gfra3^ArchT^ neurons seemed to affect behavior associated with noxious but not innocuous mechanical forces (below 1g). However, inhibition of Gfra3^ArchT^ neurons in animals with arthritis reversed sensitization associated with allodynia (below 1g stimuli) in addition to noxious pricking stimuli (above 1g stimuli) (Fig. 3c). Thus, arthritis sensitized Gfra3^ArchT^ neurons contribute to both allodynia and heightened sensitivity to painful stimuli. While the contribution of *Trpv1* and *Calca* (CGRP) expressing sensory neurons to normal pain behavior has previously been studied^42^, these genes mark several sensory neuron types including PEP1 C-nociceptors, Aδ-nociceptors as well as several types of pruriceptors and hence, the normal function of PEP1 C-nociceptors is unknown. To establish how PEP1 C-nociceptors contribute to somatosensation, we examined their role in naïve animals. In Gfra3^ChR2^ mice, subthreshold light combined with naturalistic stimuli revealed a contribution of *Gfra3*^+^ nociceptors to mechanical threshold detection, nocifensive behavior to mechanical and cold stimuli while no change was seen in heat threshold detection (Extended Data Fig. 2d). Silencing Gfra3^ArchT^ expressing C-nociceptors revealed only a small shift in mechanical withdrawal threshold (Extended Data Fig. 2d). Thus, the silencing of this neuronal type in naïve animals seems not substantially to affect pain behavioral responses but is sufficient to completely reverse arthritis pain.

**Fig. 2.**
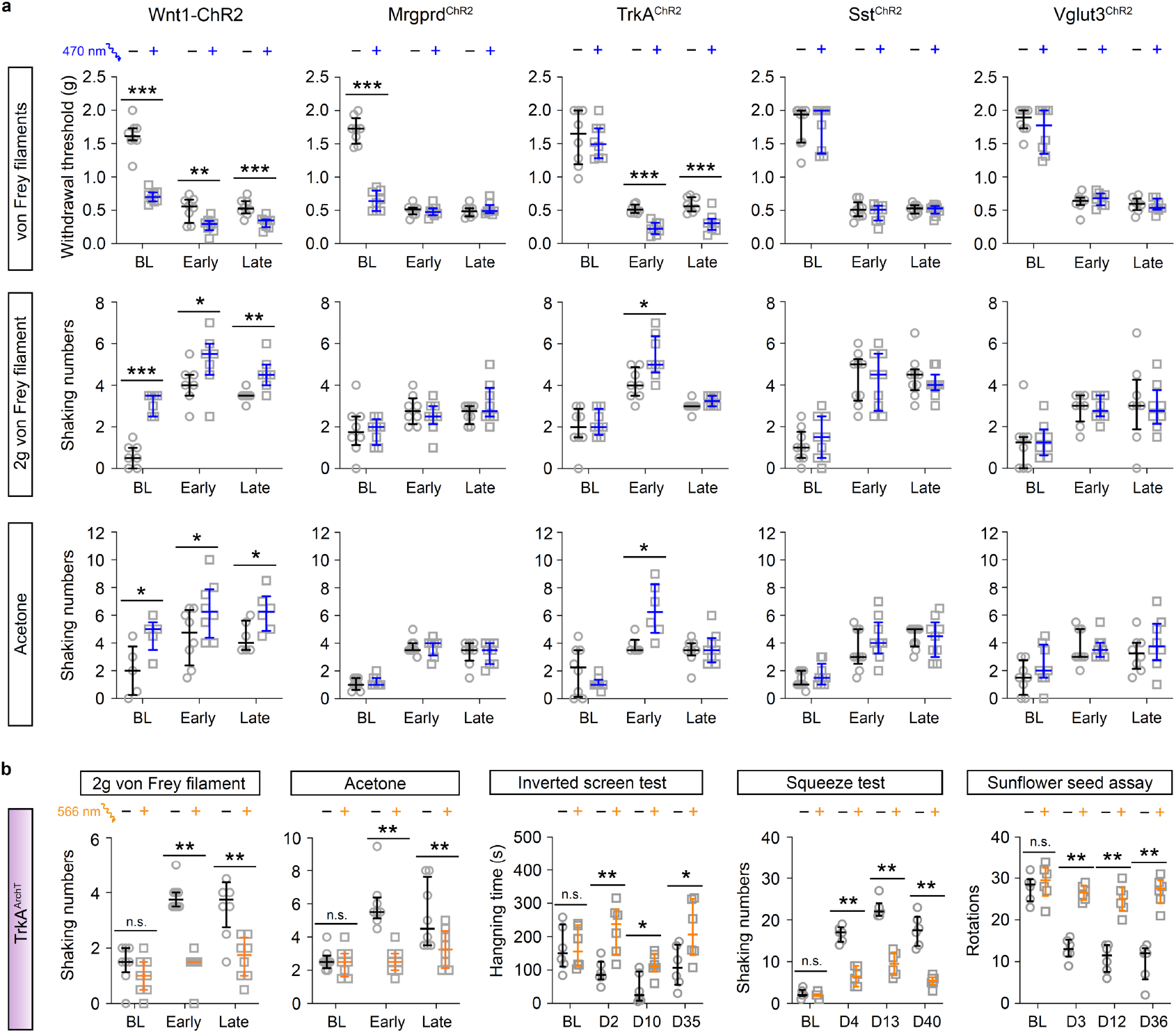
TrkA^+^ sensory neurons contribute arthritis pain. (**a**) Subthreshold photo-stimulation with blue light combined with naturalistic stimuli (mechanical threshold by von Frey filaments, mechanical pricking by 2g von Frey filament, acetone assay) prior to inducing arthritis (BL), early (during inflammation) and late arthritis (after inflammatory remission) in Wnt1-ChR2, Mrgprd^ChR2^, TrkA^ChR2^, Sst^ChR2^, and Vglut3^ChR2^ mice. Each strain contained 7-8 mice, \**p* < 0.05, \*\**p* < 0.01, \*\*\**p* < 0.001. (**b**) Mechanical/cold sensitivity and joint pain tests (including inverted screen, squeeze and sunflower seed tests) in TrkA^ArchT^ mice before and after yellow light (566 nm) application (0.1 mWatt/mm^2^, 30 min) at different timepoints after inducing arthritis (days, D, n = 8/group, \**p* < 0.05, \*\**p* < 0.01). Unpaired t test was used for inverted screen test and Mann-Whitney test was applied for other assays.

**Fig. 3.**
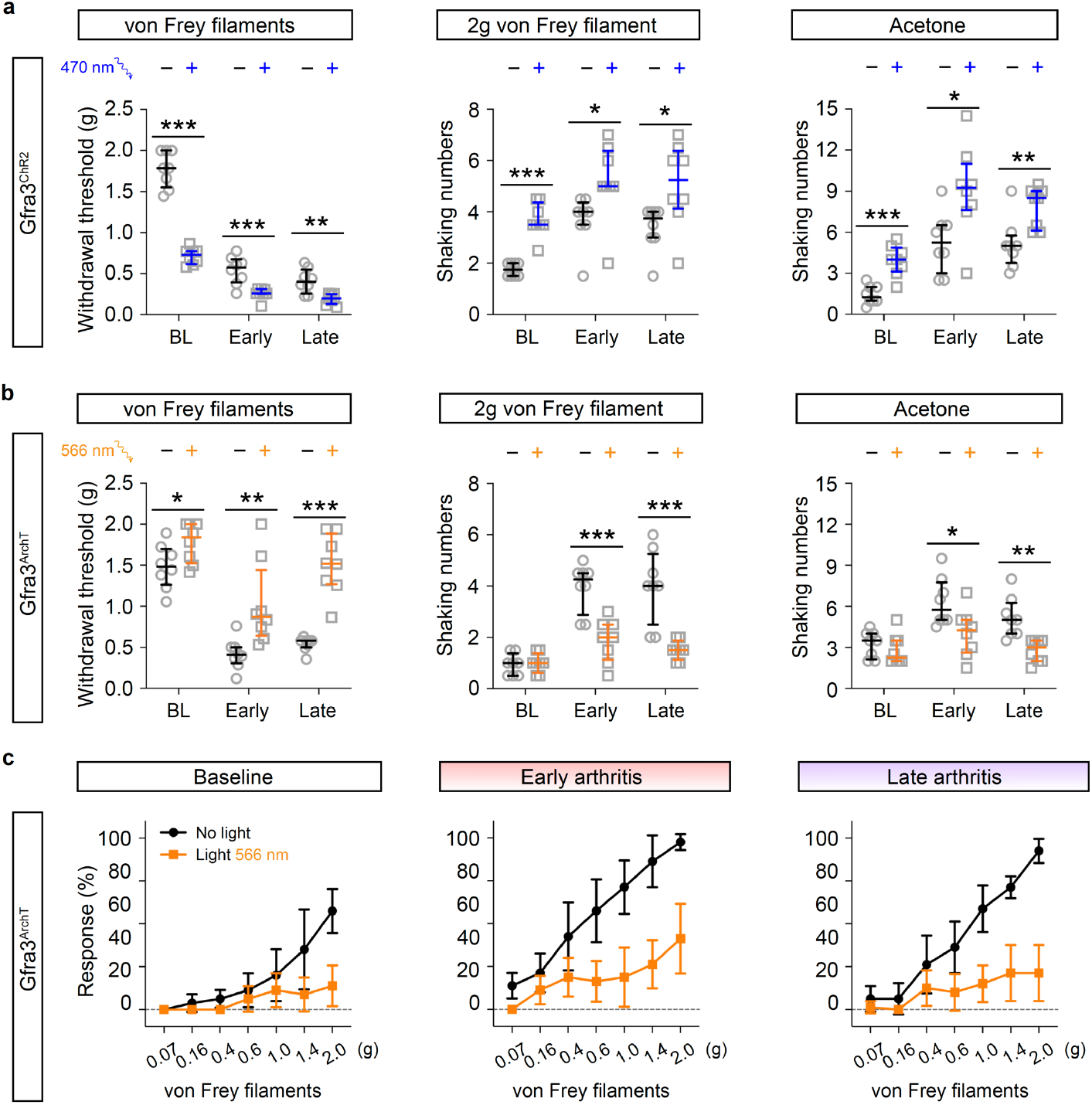
*Gfra3*^+^ C-nociceptors as a cause for pain in arthritis. (**a**) Mechanical withdrawal threshold (von Frey filaments test), nocifensive response (shaking numbers) to 2g von Frey filament and acetone test without light or with subthreshold blue light (470 nm) in Gfra3^ChR2^ mice (n = 8, MannWhitney test, \**p* < 0.05, \*\**p* < 0.01, \*\*\**p* < 0.001). BL: baseline control; early, inflammatory; late, post-inflammatory phase of arthritis. (**b**) Mechanical and cold sensitivity in Gfra3^ArchT^ mice before and after yellow light (566 nm) application (0.1 mWatt/mm^2^, 45 min) (n = 8, Mann-Whitney test, \**p* < 0.05,\*\**p* < 0.01, \*\*\**p* < 0.001). (**c**) Percentage of reflex response to different von Frey filaments without or with yellow light applied (0.1 mWatt/mm^2^, 45 min): before (Baseline) and after antibody-induced arthritis (Early and Late phases of arthritis) (n = 8).

### Type I interferons drive a transient inflammation involving a range of cell types in the DRG

Patients with RA have elevated levels of circulating cytokines^44-46^. Likewise, we observed a rapid (within an hour) increase of IL-3, IL-6, IL-10, interferon alpha (IFNα), Colony Stimulating Factor 3 (CSF3), chemokines C-C motif Chemokine Ligand 2, 3 and 4 (CCL2, CCL3 and CCL4) and C-X-C Motif Chemokine Ligand 1 (CXCL1) in serum from autoantibody injected mice which largely were normalized to control levels at 12 hours (Extended Data Fig. 3a), indicating local tissue inflammation as responsible for sensory neuron sensitization at timepoints beyond the acute phase. Furthermore, the very fast sensitization in mice with autoantibodies prior to any overt inflammation in extremities (Fig. 1b,c and Extended Data Fig. 1a) is consistent with pain independent of joint inflammation. To gain insight into the molecular underpinning of sensitization through non-neuronal and immune interactions with neurons, we scRNA-seq cells of the DRG in autoantibody injected mice.

Vehicle or arthritis inducing autoantibodies were administered and DRGs collected at 6 and 12 hours (i.e. 0.25 and 0.5 days) and day 1, 2, 12, 33 and 63 for scRNA-seq (Extended Data Fig. 3b-f and Supplementary Tables 1-4). A total of about 86 000 cells were sequenced, including nerve sheet associated cells (perineural, epineural and endoneural fibroblasts), vessel associated endothelial and lymphatic endothelial cells, pericytes, vascular smooth muscle cells, and various types of Schwann cells (satellite, non-myelinating and myelinating) as well as immune cells and neurons (Fig. 4a). All cell types identified by clustering in vehicle controls (timepoint 0) were observed at all analyzed timepoints and furthermore, we did not observe any overt alterations in proportion of cell types across timepoints after injection of autoantibodies or after onset of arthritis (Extended Data Fig. 4a,b). However, a more detailed analysis of immune cell types in the DRG led to identification of B, CD4^+^ T-helper, CD8^+^ T-cytotoxic, NK, T-reg, endoneurial macrophage, epineurial macrophage and monocyte cells (Fig. 4b) and subtypes of neurons^24^ (Fig. 4c). While the proportion of neuronal subtypes did not change (Extended Data Fig. 4c), analysis of immune cell types revealed a nearly 20-fold increase in monocytes in the DRG at 6 and 12 hours (from ∼3.5% to ∼65%) which dropped back to baseline by day 2. Simultaneously, the proportion of endoneurial macrophages decreased from ∼65% to ∼3% and back to ∼60% in the same window of 2 days (Extended Data Fig. 4d). The rapid upregulation of *Ccl2* and *Ccl4* expression in several non-neuronal and immune DRG cell types at 12h (Extended Data Fig. 4e) are consistent with the CCL-dependent recruitment of CCR2/5-expressing monocytes^47^. Macrophages exist in various configurations, ranging from inflammatory to anti-inflammatory. Unbiased identification of macrophage subtypes and assignment based on marker gene expression^48^ revealed five subtypes with a rapid and marked polarization of immune regulatory/inflammatory resolution macrophages (Mrc1-Mac, cluster 0) and activated/inflammatory macrophages (Rel-Mac, cluster 1) into proinflammatory interferon response signature macrophages (CCL2/5-Mac, cluster 2) without any change in the other two macrophage subtypes classified as antigen presenting Sparc-Mac (cluster 3) and proliferating Mki67-Mac (cluster 4) (Extended Data Fig. 4f-j). Thus, we interpreted this to represent a polarization to proinflammatory CCL2/5-Mac as well as a turnover of endoneurial macrophages from a new set of monocytes during the early stages of arthralgia (12h timepoint). Endoneural macrophages displayed a persistent alteration of a smaller set of genes throughout all the experimental timepoints (Fig. 4d and Extended Data Fig. 5a). A machine learning module trained to identify experimental versus control cells reliably identified endoneurial macrophages from mice with autoantibodies at the different timepoints up to 63 days, while epineural macrophages and monocytes were only perturbed during the first day, but not at later phases of the disease (Extended Data Fig. 5b). However, the persistent gene expression changes in macrophages were few, mostly downregulated and different from those regulated at the 12h timepoint. Top terms for KEGG pathways were related to TNF signaling pathway, indicating a sustained suppression of some proinflammatory features in endoneural macrophages during joint inflammation (12d timepoint) as well as after inflammatory remission (33d and 63d) (Extended Data Fig. 5c and Supplementary Table 5). Priming of the NOD-, LRR-and pyrin domain-containing protein 3 (NLRP3) inflammasome involves upregulated expression of NLRP3, Casp4 and IL1β through NFkB signaling^49^. Endoneural macrophages displayed a sustained downregulation of *Nlrp3* and *Casp4* at all timepoints except the first day after inducing arthritis, as well as cytokines (*Il1b*) and chemokines at later timepoints (Extended Data Fig. 5d,e), including NFkB signaling components. Differential gene expressions (Fig. 4d and Supplementary Tables 1-4) revealed a marked and sudden change within 12 hours not only in macrophages, but also in all other types of DRG cells. However, unlike macrophages, only a few gene expression alterations were observed beyond 12h in any of the other DRG cell types. The transient expansion and activation of DRG macrophages is interesting considering the essential role of resident macrophages at early, but not later stages of hypersensitivity in animals with nerve damage^50^.

**Fig. 4.**
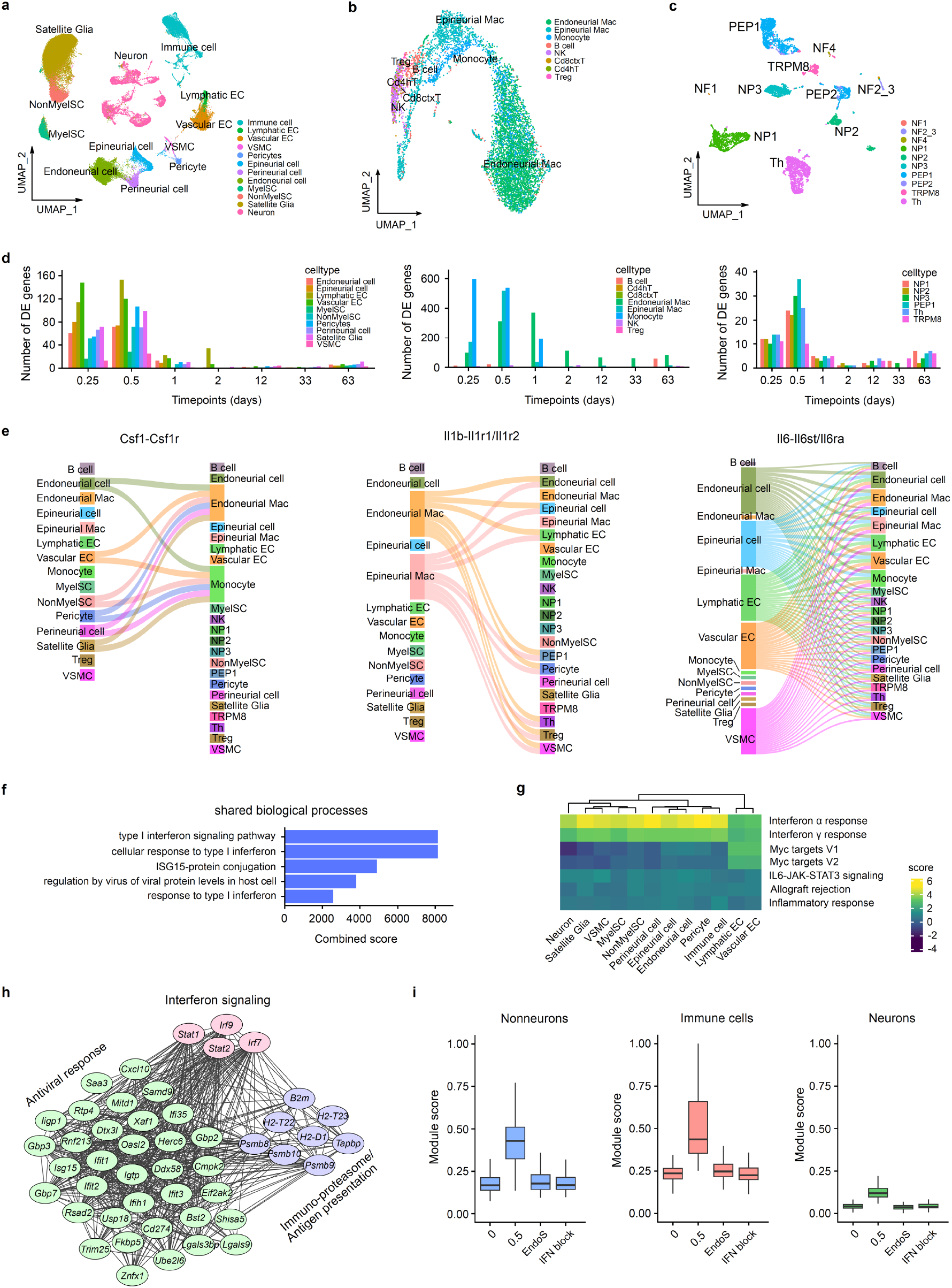
Acute cytokine storm in the DRG caused by type I interferons. (**a**) Uniform manifold approximation and projection (UMAP) showing the distribution of cell clusters (86 052 cells) from scRNA-seq of DRGs from control and arthritis mice. NonmyelSC, nonmyelinating Schwann cell; MyelSC, myelinating Schwann cell; VSMC, vascular smooth muscle cell; EC, endothelial cell. (**b**) UMAP showing the distribution of immune cells. (**c**) UMAP of neuronal clusters (6 200 cells) from scRNA-seq of DRGs from control and arthritis mice. (**d**) Number of differential expression (DE) genes of DRG cell types in arthritis mice, left graph non-neuronal cell types; middle graph, immune cell types; right graph neuronal cell types. (**e**) Ligandreceptor interaction analysis of new interactions in DRG 12h after autoantibody injection. (**f**) Combined score of pseudobulk gene ontology analyses at 12h showed relation to IFN1 signaling pathway and response to interferons. (**g**) Top gene-set enrichment analysis (GSEA) pathways at 12h datapoint. (**h**) STRING network visualization of the co-regulated modules at 12h in neurons. (**i**) Module score of different type of cells in DRG 12h after antibody injection as well as EndoS treated autoantibody injected mice and mice with arthritis preceded by antibody mediated block of the interferon receptor (IFNAR1).

Cell type assignment of neurons was unambiguous and the prediction score for identifying neuron types using machine learning was close to 1 for all neuronal subtypes analyzed, including those isolated from mice with arthritis (Extended Data Fig. 6a). This shows that arthritis does not affect the variable features defining each of the different neuron types. We next tested if the machine learning module trained to identify experimental versus control cells reliably identified sensory neurons from mice with autoantibodies. Among the sensory neuron types, NP1, NP3 and PEP1 were mostly perturbed, with the greatest perturbation at 12h (Extended Data Fig. 6b). Collectively, these results show a major molecular perturbation in sensory neurons shortly after inducing arthritis which does not involve the cell-type defining features. The marked and sudden gene expression changes after inducing arthritis led us to characterize the cellular crosstalk based on the scRNA-seq data to obtain insights into how the different DRG cell types interact with one another. Ligand-receptor interactions were inferred across the different cell types using upregulated ligand genes at the 12h peak of differentially expressed genes. Thereafter, ligand-receptor interactions were scored for enrichment of activity as compared to control mice and randomized background genes to find interactions associated with inducing arthritis versus homeostasis (Supplementary Table 6), creating an interactome of ligand-receptor interactions defining early arthritis. Notably, as visualized in Sankey plots for cytokines: *Csf1* from endoneural fibroblasts, vascular cells and satellite glia communicated with endoneural macrophages and monocytes, *Il1b* from endoand epineural macrophages had as target cells immune cells, vascular pericytes, epiand endoneural fibroblasts and neurons; *Il6* from endo- and epineural fibroblasts as well as vascular cell types interacted with all cell types of the ganglion; chemokines produced by non-neuronal and immune cells interacted mainly with immune cell types (Fig. 4e and Extended Data Fig. 6c). Thus, prior to any visible joint inflammation, unbiased analysis of cell–cell interactions identified a complex proinflammatory cellular communication network dominated by non-neuronal and immune cells of the DRG.

We used several parallel analyses strategies to obtain insights into the pathways and gene regulatory networks underlying alterations in gene expression at the 12h timepoint; gene ontology biological process of differentially expressed genes, gene-set enrichment analysis (GSEA) on hallmark genes and co-regulated gene set analysis^51^. All high scoring pathway gene sets using pseudobulk gene ontology analyses on all cells combined, or non-neuronal cells, immune cells and neurons separately were related to IFN1 signaling pathway and response to interferons (Fig. 4f and Extended Data Fig. 6d). Consistently, top GSEA pathways of non-neuronal, immune and neuronal cell types were IFNα-response although weaker significance was also obtained in most cell types for MYC, TNFα, IL6 signaling and some other hallmark gene sets (Fig. 4g and Supplementary Fig. 1). Finally, co-regulated gene expression network analyses also revealed several gene set modules of co-regulated genes related to IFN1/antiviral signaling in all cell types which were largely overlapping (Extended Data Fig. 7a,b) and, thus, were merged into one gene module with a module score representing expression of the IFN1/antiviral co-regulated genes. We used these coregulated genes for calculating the module score of pseudobulked non-neuronal cells, immune cells and neurons or of each neuronal type individually at the different timepoints after arthritis was induced in the mice. This revealed a rapid and transient expression of antiviral genes in all the different DRG cell types (Extended Data Fig. 7c,d). Among the genes were interferon signaling molecules, immunoproteasome/antigen presentation and inhibitors of viral propagation (Fig. 4h, neurons).

Activation of the IFN1 receptor (IFNAR1/IFNAR2 heterodimer) leads to STAT1 and STAT2 phosphorylation and recruitment of IRF9, these together forming the IFN-stimulated gene factor 3 complex (ISGF3) that activates interferon stimulated genes (ISGs)^52^. We used SCENIC to identify gene regulatory networks created from master transcription factors and their gene targets (i.e. regulons) in all DRG cell types at all timepoints (Extended Data Fig. 8a and Supplementary Table 7)^53^. We found a total of 27 regulons, of which 4 were associated with autoantibody injection. These were increased during the first 24h and included IRF7, IRF9, STAT1, STAT2 (n = 42-66 genes). In immune cell types IRF7, IRF8, STAT1, STAT2 (40-450 genes) were the most increased and in non-neuronal cell types IRF7, IRF9, STAT1, STAT2 (18-242 genes). Certain non-neuronal cells also had gene regulatory networks downstream of *Cebpd*. CEBPD is involved in immune and inflammatory response genes and suggested to be activated by IL-1β, TNFα and IL-6 and transactivates cytokine expression^54^, consistent with non-neuronal cell types and macrophages as the main cellular source producing pro-inflammatory mediators in the DRG at early phases of disease as identified by ligand-receptor analyses in Fig. 4e and Extended Data Fig. 6c.

Taken together, the above results implied IFN1 as the major mediator driving transcriptional alterations in dorsal root ganglion cells associated with autoantibody-induced arthralgia. The results suggest an induction of ISGs through interferon α/β activation of type I interferon receptors rather than through cytosolic sensors and pathways or toll-like receptors. However, to exclude that the ISGs were triggered by DNA, RNA or pathogen contaminants in the autoantibody mix used to induce arthritis we cleaved the Fc-glycan needed for antibody effector function through binding to Fc-receptors, using the endoglycosidase from the human pathogen *Streptococcus pyogenes*, EndoS^55^. We treated the cocktail of cartilage-binding antibodies with EndoS, repurified the antibodies and thereafter administered them to the mice at the same concentration as used in the previous experiments. Animals administered EndoS treated autoantibodies did not develop allodynia or hyperalgesia (Extended Data Fig. 8b). DRGs from EndoS treated and control mice were collected and scRNA-seq at the 12h peak timepoint for ISGs expression. Clustering revealed all cell types previously identified in the DRG to be present in both control and experimental mice, including nerve sheath associated cells, vessel associated cell types, various types of Schwann cell glia, neurons and immune cells (Extended Data Fig. 8c,d), including subtypes of immune cells (Extended Data Fig. 8e). Mice administered EndoS treated autoantibodies did not show any induction of ISG module and had no monocyte infiltration (Fig. 4i and Extended Data Fig. 8e), nor were any differentially expressed genes identified between control and experimental mice (Extended Data Fig. 8f). This shows that gene regulation as well as hyperalgesia is caused by functional autoantibodies. To establish whether the gene expression changes can be directly ascribed to type I interferons, we administered an IFNAR1 inhibiting antibody (MAR1-5A3) and 1 hour later administered the arthritis autoantibodies. DRGs were isolated for scRNA-seq at the peak of differential gene expression (12h). Like the EndoS experiment, mice with IFN block displayed all previously identified cell types in the DRG, had an absence of monocyte infiltration, ISGs, absence of regulons associated with arthritis, as well as a lack of any differentially expressed genes (Fig. 4i and Extended Data Fig. 8c-f). These results show that the cartilage binding antibodies, relying on Fc-glycan needed for local immune complex formation induce IFN1 release and activation of interferon receptors^56^, resulting in ISG expression. Furthermore, blocking IFN1 leads to a complete reversal of the inflammatory status of the DRG in the early phase, uncovering IFNs as the dominant proinflammatory factor driving inflammation in the ganglion.

### A persistent alternative interferon signaling pathway in arthritis causes sensory neurons hyperexcitability

The dominant role of interferons at early periods after the injection of autoantibodies made us consider a role also during and after the active period of arthritis. We examined whether IFNs were continuously produced and induced intracellular signaling in neurons and by this contribute to sensitization and pain in arthritis. Because the rise in systemic type I IFN levels appeared in the early phase and was transient in our mouse model and likewise in patients,^57, 58^ a persistent IFN-dependent neuronal hyperexcitability likely arises from local IFN1 produced in the tissue. Because IFN1 transcripts are not polyadenylated, we were unable to identify expression in the scRNA-seq data and instead performed quantitative PCR of DRGs for all 14 *Ifna* genes combined and for *Ifnb* in mice with experimental rheumatoid arthritis. Autoantibody-induced arthritis led to a sustained local transcription of IFNs in the DRG throughout all timepoints of the disease (Fig. 5a). RNA *in situ* hybridization with probes designed at conserved regions of *Ifna* genes revealed primarily neuronal expression while *Ifnar1* was broadly expressed in the DRG including *Calca*^+^ neurons (Fig. 5b), as well as endothelial and immune cells as revealed by scRNA-seq (Fig. 5c). To assess whether IFN1 can alter excitability of sensory neurons, we used patch-clamp electrophysiology. Small to medium size mouse DRG neurons exposed to IFNα3 for one hour were markedly sensitized as seen by a lower threshold for activation, increased action potential firing and a shorter latency to first spike (Fig. 5e,f and Extended Data Fig. 9a), consistent with a previous study^16^, suggesting increased intrinsic excitability as a mechanism for sensory dysfunction in arthritis in our mouse model.

**Fig. 5.**
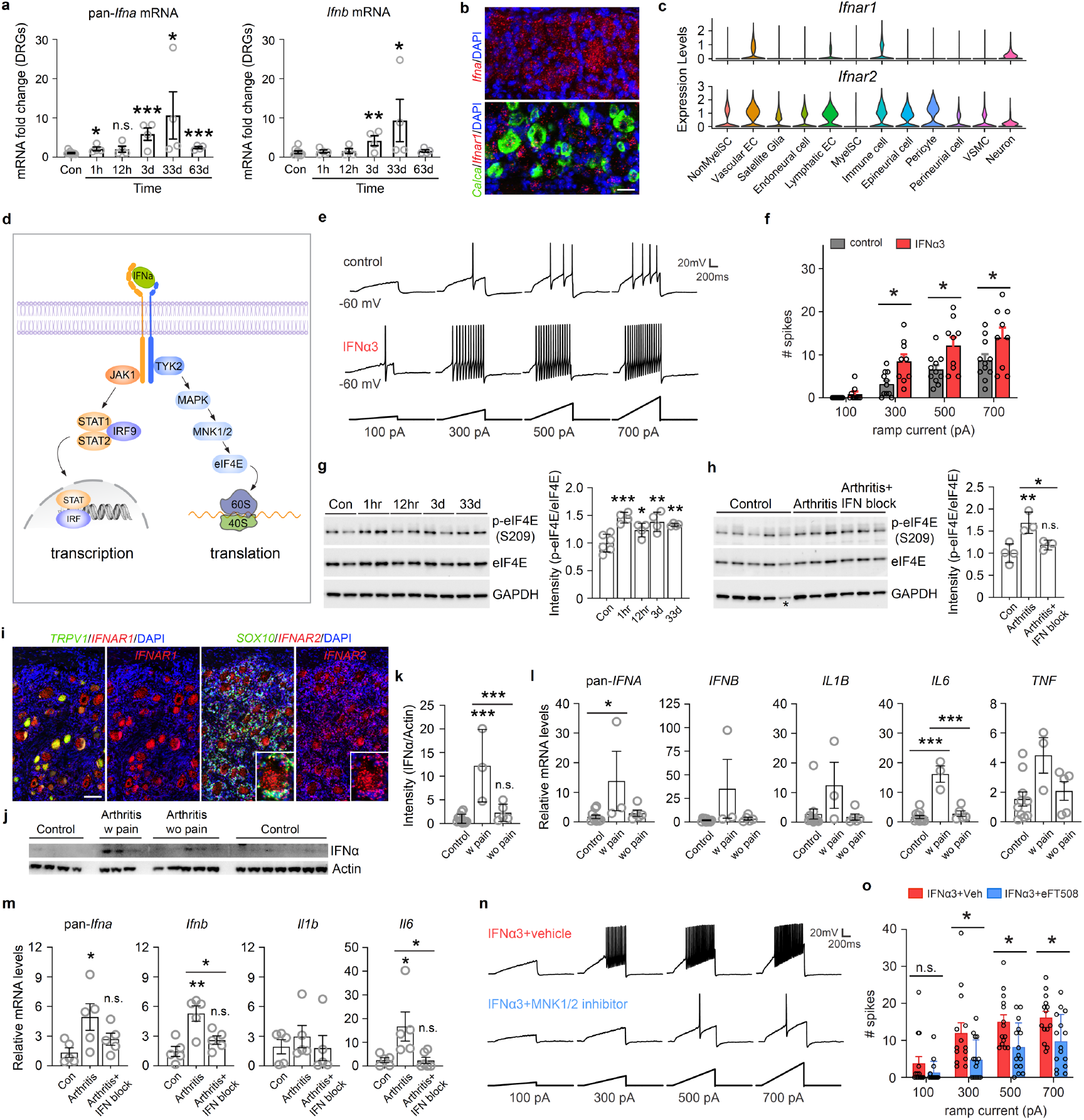
Sustained interferon signaling drives arthritis pain. (**a**) Expression changes of *Ifna* and *Ifnb* mRNA levels at different times (hours and days) after autoantibody injection in mouse DRGs by qPCR. Pan-*Ifna* primers were used for all *Ifna* genes. Each time point of arthritis contains 4 mice, whereas control group contains 9 mice, \**p* < 0.05, \*\**p* < 0.01, \*\*\**p* < 0.001. (**b**) Representative RNAscope images of *Ifna* mRNA (top) and *Ifnar1* mRNA (bottom) expression in control mouse DRGs. Scale bar = 25 µm. (**c**) Expression levels of type I IFN receptors *Ifnar1* and *Ifnar2* in different cell types from mouse DRG scRNA-seq dataset. (**d**) Illustration of transcriptional and translational pathways activated by type I interferons. (**e**) Current-clamp responses of cultured mouse DRG neurons to IFNα3 during different current ramp injections. (**f**) Plots of number of spikes recorded from control and IFNα3 stimulated DRG neurons under different ramp currents (control, n = 11; IFNα3, n = 9, \**p* < 0.05). (**g**) Phosphorylation of eIF4E (Ser209) (normalized to eIF4E) in mouse DRGs by western blotting at different times after antibody injection. Each time point contains 4 mice, except control group (before injection), which includes 6 mice, \**p* < 0.05, \*\**p* < 0.01, \*\*\**p* < 0.001. (**h**) Effect on phosphorylation of eIF4E by i.p. injection of IFNAR1 blocking antibody (IFN block) in arthritis mice (con, control mice, n = 4 in each group, \**p* < 0.05, \*\**p* < 0.01). The fifth control sample labeled with an asterisk on GAPDH band was not included for quantification. (**i**) Representative RNAscope images of human DRGs from female healthy donors showing *IFNAR1* mRNA (*TRPV1*/*IFNAR1*, left) and *IFNAR2* mRNA (*SOX10*/*IFNAR2*, right) expression in neurons. Inset for *IFNAR2* on the right showed co-localization of *SOX10* and *IFNAR2* also in satellite glia cells. Scale bar = 100 µm. (**j**) Western blot analysis shows IFNα expression in human DRGs from healthy controls (n = 11), RA patients with pain (Arthritis w pain, n = 3) and RA patients without pain (Arthritis wo pain, n = 5). (**k**) Quantification of IFNα expression by western blotting in (**j**) within healthy control, RA patients with pain (w pain) and RA patients without pain (wo pain), \*\*\**p* < 0.001. (**l**) mRNA expression fold changes of *IFNA, IFNB, IL1B, IL6* and *TNF* in human DRGs (\**p* < 0.05, \*\*\**p* < 0.001). (**m**) mRNA expression fold changes of *Ifna, Ifnb, Il1b* and *Il6* in mouse DRGs from control (n = 5), arthritis (n = 5) and arthritis mice treated with IFNAR1 blocking antibody (IFN block, n = 5), *p < 0.05, **p < 0.01. (**n**) Action potential responses in the current clamp setting following current injections of cultured mouse DRG neurons exposed to IFNα3 with or without MNK1/2 inhibitor eFT508. (**o**) Plots of number of spikes recorded from IFNα3 stimulated DRG neurons in presence or absence of MNK1/2 inhibitor under different ramp currents (IFNα3 + Vehicle, n = 14; IFNα3 + eFT508, n = 15, \**p* < 0.05). qPCR and western blotting data were analyzed with one-way ANOVA followed by Turkey’s multiple comparisons test, whereas patch clamp analysis was performed with two-way ANOVA followed by Šídák’s multiple comparisons test.

Although most interferon-induced gene expression alterations are produced on demand by ISGF3, the response is transient and resolved through feed-back inhibition^59, 60^. Thus, the lack of interferon stimulated gene expression at inflammatory and post-inflammatory phases of hyperalgesia and arthralgia despite a continuous presence of IFN1 in the DRG indicated an alternative persistent signaling as a plausible cause for pain in arthritis. Alternative “non-ISGF3” antiviral signaling pathways which does not lead to gene expression alterations include mitogen-activated protein kinase (MAPK)-integrating kinase (MNK) 1/2 and eukaryotic translation initiation factor 4E (eIF4E), which modulates translation^61^ (Fig. 5d). To obtain evidence for persistent interferon signaling through the phospho-serine-threonine MNK1/2-eIF4E pathway in vivo in arthritis, we collected DRGs from mice at 1h, 12h, 3d and 33d after induced arthritis and compared to mice without arthritis (control). eIF4E (Ser209) phosphorylation can be used as a direct measure of MNK1/2 activity, since they are the only kinases that phosphorylate eIF4E^62^. Phospho-eIF4E (S209) was significantly elevated in mice with arthritis at all timepoints (Fig. 5g and Supplementary Fig. 2a). The increased eIF4E (Ser209) was caused by IFN1, because inhibition of ligand interaction with its receptor in vivo by administration of the IFNAR1 blocking antibody on day 33 after inducing arthritis reversed the increase of eIF4E (Ser209) phosphorylation (Fig. 5h and Supplementary Fig. 2b). Thus, engagement of IFN receptors by IFN1 leads to persistent signaling through MNK1/2-eIF4E in DRG in mice with arthritis.

*In situ* hybridization for interferon receptor expression in human DRG showed *IFNAR1* expression in both *TRPV1*^+^ nociceptors as well as *TRPV1*^-^ sensory neurons, while *IFNAR2* receptor was likewise expressed in most or all sensory neurons, it was also expressed in non-neuronal cells such as *SOX10*^+^ satellite glia that cover the surface of the neuronal cell bodies (Fig. 5i and Supplementary Fig. 3). Thus, human sensory neurons carry receptors necessary to respond to the presence of IFN1. To test whether our results obtained in the mouse arthritis model translate into patients with RA, we analyzed human DRG from patients with RA and joint pain, patients with RA but without joint pain and controls (Supplementary Table 8). Western blot analysis of IFNα protein levels revealed increased protein expression in the DRG of individuals with RA and pain as compared to the other groups (Fig. 5j,k). Furthermore, quantitative PCR for all 13 IFNα genes using primers targeting conserved regions and for IFNα revealed increased mRNA expression in DRG from patients with RA and joint pain as compared to those without pain or control group (Fig. 5l). Thus, these results evidence an increased expression of IFN1 in the DRG of the mouse autoantibody model as well as in individuals with RA and pain, while those with RA without pain were like individuals without RA. In addition, we found that several other cytokines were elevated in human DRG of patient donors with RA and pain, most notably IL-6 (Fig. 5l). When re-examining autoantibody-induced arthritis mice on day 33, although expressed at overall very low levels, we confirmed by quantitative PCR increased IL-6 levels in addition to IFN1 also in the mouse model (Fig. 5m). IL-6, like IFN1, signal through the eIF4E complex and sensitizes sensory neurons^63, 64^. Thus, both IFN1 and IL-6 could contribute to alterations in sensory neuron excitability by converging onto MNK1/2. We therefore examined whether elevation of IL-6, along with IFNα and IFNβ, in DRG of mice with autoantibody-induced arthritis relied on interferon receptor signaling using the IFNAR1 blocking antibody. Inhibiting the interferon receptor on day 33 in mice with arthritis reversed the increase of *Il6* along with *Ifna* and *Ifnb* (Fig. 5m). We performed patch-clamp recordings on small size mouse DRG neurons under perfusion with IFNα3 in the presence or absence of the MNK1/2 inhibitor eFT508. Inhibiting MNK1/2 reversed IFNα3-induced hyperexcitability by normalizing the threshold for activation and preventing IFNα3-induced increased action potential firing (Fig. 5n,o and Extended Data Fig. 9b). These results suggest that type I IFNs and possibly also IL-6 induced by IFN1 in vivo is engaging MNK1/2 and that this is causative for sensory neuron hyperexcitability.

### Reversal of pain by inhibition of the IFN1-MNK1/2-eIF4E signaling pathway

If the observed neuronal hyperexcitability by the IFN1-MNK1/2-eIF4E pathway is the cause for pain in arthritis, pathway inhibition in vivo should reverse that pain. To directly establish if IFN1 and the persistent IFN1 signaling pathway are responsible for pain in arthritis, allodynia and pain behavior was assessed in arthritis mice administered with the IFNAR1 blocking antibody. Blocking interferon signaling prior to administering the autoantibodies prevented onset of allodynia and noxious mechanical hyperalgesia. However, the effect was transient and by 2.5 days after administration of the IFNAR1 antibody the effects were reversing and by 3.5 days mechanical allodynia and hyperalgesia were similar to untreated animals injected with autoantibodies (Extended Data Fig. 10a), consistent with previous studies dosing every other day to maintain signaling inhibition^65^. Inhibition did not affect the disease inflammatory score (Extended Data Fig. 10b). Blocking with the IFNAR1 blocking antibody in mice with already established arthritis at day 22 and after resolution of inflammation at day 45 reversed mechanical allodynia and mechanical hyperalgesia for 2 days (Extended Data Fig. 10a). Active RA is characterized by an involvement of several different cytokines elevated both in circulation and in peripheral tissues, such as the joint synovium^66^, which leads to joint inflammation, vascularization and joint swelling. When examining whether inhibition of IFN receptor activation can ameliorate nociceptive hypersensitivity at the peak of swelling, we found an efficient reversal of both mechanical allodynia and hyperalgesia (day 12 of arthritis, Extended Data Fig. 10c). Notably, joint pain assessed by the paw squeeze test, deficits of forepaw dexterity in the sunflower seed assay and limb usage disability assessed in the inverted screen test were also reversed at the post-inflammatory chronic phase (Fig. 6a). IFN1 receptors are associated with tyrosine kinase 2 (TYK2) required for receptor phosphorylation and receptor-induced signal transduction^67^. To test if inhibition of TYK2 like inhibition of the interferon receptor can reverse arthritis pain-behavior, we used the allosteric TYK2 inhibitor deucravacitinib, used to treat psoriasis^68^. Deucravacitinib administered daily for 9 days potently reversed allodynia, hyperalgesia, joint pain, and improved dexterity and limb function with an effect already after the first day (Fig. 6b and Extended Data Fig. 10d). Inhibiting MNK1/2 using the eFT508 inhibitor as well as its substrate eIF4E using the 4EGI inhibitor completely reversed allodynia, hyperalgesia, joint pain, and improved dexterity and limb usage in mice with arthritis (Fig. 6c,d and Extended Data Fig. 10e,f). IFNα was sufficient by itself to cause sensitization through MNK1/2, since local cutaneous IFNα3 injection in mice led to allodynia and hyperalgesia which were reversed by the MNK1/2 inhibitor (Extended Data Fig. 10g). Collectively, these data reveal that sustained IFN1 signaling engages the MNK1/2-eIF4E pathway which is the cause for increased excitability in nociceptive neurons and for joint pain.

**Fig. 6.**
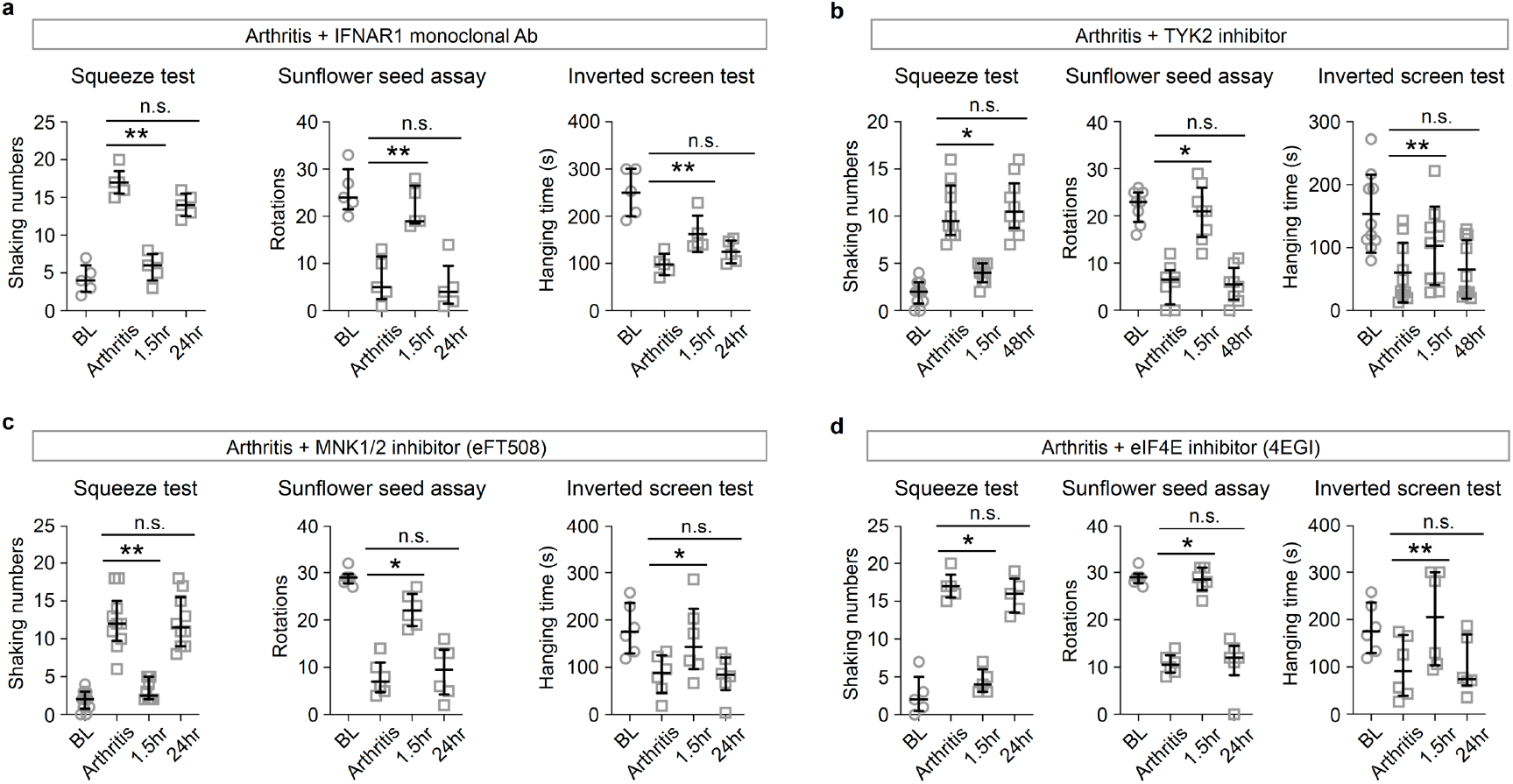
Inhibition of interferon signaling restores limb function and reverses pain in arthritis. (**a**) Systemic administration of IFNAR1 mAb (i.p., 40 mg/kg) reversed chronic joint pain, impairment of dexterity and limb function in the squeeze test, sunflower seed assay and inverted screen test (C57BL/6N, n = 5, \*\**p* < 0.01). (**b**) Oral administration of TYK2 inhibitor (15 mg/kg) similarly reversed sensory deficits associated with arthritis (n = 8, \**p* < 0.05, \*\**p* < 0.01). (**c**) MNK1/2 inhibitor (i.p., eFT508, 1 mg/kg) reversed sensory deficits associated with arthritis (n = 10, \**p* < 0.05, \*\**p* < 0.01). (**d**) eIF4E inhibitor (i.p., 4EGI, 15 mg/kg) reversed sensory deficits associated with arthritis (n = 6, \**p* < 0.05, \*\**p* < 0.01). In all graphs show behavior prior to inducing arthritis (BL), at the post-inflammatory phase following antibody-induced arthritis (Arthritis), post inflammatory phase of arthritis 1.5h after administration of compounds (1.5hr) and after washout of compounds (24hr or 48hr). Squeeze test and sunflower seed assay were analyzed with Fridman test followed by Dunn’s multiple comparisons test. The inverted screen test was performed with one-way ANOVA followed by Šídák’s multiple comparisons test.

## Discussion

Identifying the neuronal types responsible and the causative mechanism is fundamental for understanding pain in arthritis. Here, we used mouse genetics, optogenetics, large scale transcriptomic measurements, computational biology, mouse pain and dexterity behavior, physiology, and biochemistry to chart a molecular atlas of neuro-immune interactions and molecular mechanisms of pain in arthritis across disease progression. We provide evidence for acute and persistent inflammatory phases in the DRG. In the acute phase, exposure to autoantibodies engaged immune, non-neuronal and neuronal cell types of the DRG and numerous circulating and locally produced cytokines and chemokines lead to inflammatory gene transcriptional networks in all cell types of the ganglion. In the persistent phase, inflammatory expression of cytokines and chemokines largely subsided, and gene regulatory changes were absent except for alterations consistent with the presence of anti-inflammatory macrophages. We provide evidence that: 1) Pain in arthritis is conveyed through a discreet and molecularly defined neuronal type; 2) Acute inflammation in the DRG as well as persistent pain in arthritis is associated with elevated local IFN1 levels, and hence, the mechanism of pain is largely unrelated to that of arthritis disease activity; 3) Interferon signaling in the acute phase involves antiviral ISGs while a persistent alternative signaling pathway drives pain in arthritis; 4) The persistent MNK1/2-eIF4E interferon signaling results in hyperexcitability of sensory neurons, sensitization and pain; 5) Interferon and MNK-eIF4E signaling inhibition effectively alleviate joint pain, restores paw function and dexterity in mice; and 6) In humans with RA and pain, local interferon production in the DRG is elevated, like in mouse model and hence, could be causative for pain in RA.

### The sensory neurons of pain in arthritis

Based on our results, cartilage autoantibody-induced arthritis in mice leads to several sensory related dysfunctions, including arthralgia, allodynia, cold hyperalgesia, painful and swollen joints and reduced dexterity and disability of paw and limb function. Because there has historically been little attention on quantitatively assessing hand functions and dexterity in patients with RA, there is insufficient insight into deficits beyond inflammation and pain. Nevertheless, the phenotype observed in our mouse model appears to correlate with what is known based on recent studies^69^. Previous animal studies on pain in arthritis have focused on measuring cutaneous nociception. We find that cutaneous sensitization correlates with joint pain and loss of dexterous paw usage and therefore is an acceptable proxy for studying sensory dysfunction in animal models of arthritis. Our initial idea was to infer the cell type responsible for pain in arthritis by identifying the sensitized sensory neuron type but found that all nociceptor types were sensitized. Consistently, all nociceptors contained ISG expression and interferon induced gene regulatory networks. IFN1 and ISG gene expression signature are mostly present in blood in the preclinical phase while in patients with RA it is present in the synovium^70^. However, its presence in the DRG has not been assessed. We found a persistent elevation of IFN1 within the DRG in arthritis of both mice and human. Thus, neuronal excitability and pain may be caused by the presence of interferons at any or all of these sites. The current view of some of the symptoms of RA such as stiffness, reduced grip strength and dexterous use of hands is associated to the severity of inflammation and hence, a consequence of joint inflammation and swelling^69^. Our results suggest that this idea is incomplete, and that stiffness and fine motor functions is due to sensory-motor dysfunction caused by interferons across different populations of sensory neurons, because inhibiting interferon signaling reversed not only pain behavior, but also dexterity and overall paw function seemingly independent of disease progression and remission. Therefore, interferon-induced alterations of sensory transduction and transmission in neuronal types beyond those causing joint pain is possible, an idea consistent with sensitization of myelinated joint innervating low-threshold mechanoreceptors in experimental arthritis^71^. However, since our study was focused on the main clinical problem in RA, joint pain, we did not specifically study this aspect beyond ascertaining interferon-induced gene expression changes in touch and proprioceptive neurons and the behavioral benefits of interferon inhibition therapy on dexterity and paw usage.

Nociceptive sensory afferents are highly diverse and differentially contribute to protective reflex withdrawal and the perception of pain and furthermore are tuned to respond to cold, heat and mechanical nociception. It is possible that different sensory neuron types also are responsible for pain phenotypes such as allodynia, burning, pricking/stinging and aching pain^42^. Although sensory neuron types have been molecularly identified and classified^42^, the populations of neurons responsible for pain in arthritis remained unknown beyond identification of both Aand C-fiber nerves in the joints, often containing neuropeptides such as CGRP (*Calca* gene) and the thermal transducer TRPV1^28, 29^. The CGRP^+^ and TRPV1^+^ neurons have long been known to be very heterogenous and include C- and A-fiber nociceptors detecting heat, cold and mechanical stimuli as well as pruritogens^72-75^. Unlike studies inhibiting neural activity of CGRP^+^ and TRPV1^+^ neurons^42^, inhibition in the present study of the C-fiber GFRA3^+^ neurons, which represent a subpopulation of CGRP^+^ and TRPV1^+^ neurons, unexpectedly did not lead to any marked deficit of normal pain sensitivity to mechanical, heat and cold yet completely reversed pain in arthritis. There is a linear relationship between an increase in C-nociceptor impulses and the magnitude of reported pain rating in humans^76^. C-nociceptor input can increase and contribute to hyperalgesia also by recruitment of so-called silent nociceptors (also called mechanically insensitive afferents). These are normally relatively insensitive to mechanical stimuli in the absence of tissue injury but become responsive when sensitized by inflammation^77^. Consistent with a contribution to pain in arthritis, silent nociceptors are found mainly in deep tissues such as the knee joint, and application of inflammatory compounds leads to spontaneous activity as well as increased response to gentle stimulation which has led to the suggestion that pain in inflamed tissue results from sensitized silent nociceptors^71^. In mouse, *Gfra3* is expressed in the silent nociceptors that acquires mechanosensitivity when exposed to the inflammatory mediator nerve growth factor^42, 78^. The ability of inhibition of *Gfra3*^+^ neurons to reverse pain in arthritis without much effect on normal pain sensitivity in our experiments is consistent with a recruitment of silent nociceptors normally not involved in basal pain transduction.

### Acute cytokine storm and sustained IFN1 signaling in the DRG

RA characterized by systemic inflammation eventually leading to synovitis. Aligning with our results in the mouse model, both autoantibodies^79-81^ and inflammatory threshold^82, 83^ precedes onset of arthritis^44, 84, 85^, and further maturation or switch of specificity of the autoantibody response are believed to be important in triggering the transition from systemic immunity to synovitis in patients who progress to rheumatoid arthritis. Activation of arthritogenic autoantibodies with competence of Fc-gamma receptor binding led to DRG inflammation even before any overt joint inflammation was observed. We found a fast and sudden increase of inflammation gene expression signatures in the DRG which circulating cytokines also could contribute to setting into motion. It was characterized by an inflammation signature in all cells in the DRG, including neurons, glia, vascular cells, immune cells, and cells of the endo- and perineurium. The broad DRG inflammatory response was also manifested by increased expression within the DRG of multiple cytokines, marked appearance of numerous inflammatory ligand-receptor interactions across the DRG cell types and consistently, a manifestation of down-stream gene-regulatory networks in immune, non-neuronal and neuronal cells of the DRG. However, the most robust gene-regulatory network in all cell types of the DRG was the antiviral interferon-stimulated gene expression. Furthermore, in contrast to immune and non-neuronal cells, neurons contained mainly one gene-regulatory network, namely that induced by IFN1. Consistently, at the top of the hierarchy of the marked DRG inflammatory response was IFN1 since inhibition prevented all gene expression changes in all cell types of the ganglion. Furthermore, IFN1 was continuously produced within the DRG itself in the prodromal, inflammatory, and post-inflammatory phases. Thus, autoantibodies forming local immune complexes trigger the production and release of IFN1. The importance of ongoing IFN1 signaling is consistent with the finding that preventive therapy did not have a sustained effect and established chronic pain could be overcome by inhibiting ongoing IFN1 receptor signaling both during inflammation and after remission. Thus, our results suggest that unlike synovitis, where IFN1 seems to play an inferior role in inflammation, they are central for pain and that remaining pain that persists after control of inflammation is caused by a local subclinical inflammatory response. IFN1 interacted directly with sensory neurons and lowered the threshold of activation, increased action potential firing, sensitized the nerves, and caused allodynia, hyperalgesia and joint pain. Thus, our results are consistent with that IFN1-induced hyperexcitability of *Gfra3*^+^ nociceptors drive pain at all phases of the disease.

### A persistent interferon-induced MNK1/2-eIF4E pathway activation as a cause for sensory plasticity

The innate antiviral immune system is one of the first lines of cellular defense against viral pathogens. Interferon-induced gene expression through ISGF3 in neurons included expected effectors such as virus translation and replication inhibition proteins belonging to the “IFN-induced proteins with tetratricopeptide repeats” IFIT1, IFIT3 and UBE2L6, TRIM25, USP18, inhibition of virus budding such as RSAD2, BST2^53^, and immunoproteasome/antigen presentation (Psmb8,9,10 and MHC class I genes). However, we did not consider that these genes caused hyperalgesia and pain in arthritis since the response was transient and had completely subsided at a time when sensitization and increased pain behavior continued in the phases of both active arthritis and post-arthritis. This conclusion is supported by data showing that ISG expression by STING activation exerts antinociceptive effects^86-88^. While persistent IFN1 signaling is known to contribute to malfunction and disease activity in systemic autoimmune diseases it is primarily believed to be conspicuously controlled at the gene transcription level^59, 89^. Nevertheless, alternative “non-ISG” antiviral signaling pathways driving translational changes restricting viral propagation are believed to act in concert with gene transcription, including an activation of MNK1/2 and eIF4E which controls cap-dependent translation^16, 61, 90^. IFN1, along with several other cytokines and growth factors can activate this alternative pathway^90, 91^ and have been found to regulate pain plasticity^16, 91, 92^, including the suppression of ectopic activity in human sensory neurons^93^. Thus, while multiple negative feedback mechanisms suppressing excessive signaling as well as resolving the antiviral response mediated through JAK/STAT signaling are known^59, 60^, and explains the transient nature of the interferon-stimulated gene expression in sensory neurons, we found that interferon receptors still accessed the MNK1/2-eIF4E signaling pathway in arthritis. Activation of the MNK1/2-eIF4E pathway and pain relied on IFN1 interaction with its receptor, MNK1/2 and eIF4E. Cap-dependent translation limits protein synthesis in eukaryotic cells. The binding of eIF4E to the mRNA cap structure initiates ribosome recruitment and as such regulate the synthesis of a subset of proteins by controlling mRNA translation initiation^94^. The complex and unresolved nature of translational regulation by this pathway in sensory neurons which could explain the increased excitability in arthritis^95-97^, precludes at this point a targeting of effector proteins of hyperexcitability.

### Pain relief in RA

Due to a lack of effective treatment options for pain, patients often resort to analgesics with limited efficacy that do not target the cause of chronic pain. However, it was recently found that targeting multiple cytokines with the broad-spectrum cytokine inhibitor baricitinib (JAK1/2 inhibitor) lessens pain associated with RA, as demonstrated in the Phase clinical trial RA-BEAM (NCT01710358). As JAK1 is required for interferon receptor activation, it is anticipated to also block interferon signaling. However, baricitinib increases risk for major cardiovascular problems such as blood clots, heart attack, stroke, cancer, immune system problems and death which limits utility. Beyond a broad JAK inhibition there is limited advance of treatments for pain in arthritis. Our results inhibiting the persistent interferon signaling pathway distinguish itself from current strategies in that it targets only the causative mechanism for pain in arthritis. The approach of treating pain in arthritis with compounds inhibiting the IFN1 signaling pathway has been extensively validated in our autoantibody induced arthritis mouse model and its clinical relevance is indicated by expression of the interferon receptors in human sensory neurons and the marked elevation of IFN1 in the DRG of patients with RA and pain. The identified interferon pathway includes effector proteins with already approved, or drugs in clinical trial, for other indications (e.g. anifrolumab deucravacitinib, tomivosertib) and expected side effects are largely known and limited. This should present a significant advantage for clinical translation of our findings. We believe that our discovery could represent a new approach towards treating pain in arthritis.

## Materials and Methods

### Animals

Wild type C57BL/6N mice (adult, 8-9 wk) were ordered from Charles River (Scanbur AB). *Wnt1-Cre* (*Wnt1-Cre*, JAX #003829), *Slc17a8*^*cre*^ (*Vglut3*^*Cre*^, JAX #028534), *Gfra3*^*cre/ERT2*^ (*Gfra3*^*CreERT2*^, JAX #029498), *ROSA26*^*Tomato*^ (Ai14, JAX #007914), *ROSA26*^*ChR2-EYFP*^ (*ROSA26*^*ChR2*^, Ai32, JAX #012569) and *ROSA26*^*ArchT-EGFP*^ (*ROSA26*^*ArchT*^, Ai40D, JAX #021188) were ordered from The Jackson Laboratory. *Sst*^*cre*^ was a generous gift from Jens Hjerling-Leffler (*Sst*^*Cre*^, JAX #013044). *Mrgprd*^*cre*^ was ordered from Mutant Mouse Resource & Research Centers (*Mrgprd*^*Cre*^, MMRRC_036118) and *Ntrk1*^*cre/ERT2*^ (*TrkA*^*Cre-ERT2*^) mice were generated in the lab as previously described^39^. All the strains were back crossed to C57BL/6N wildtype mice at least for 3 generations before used for breeding. The following crosses were done: *Wnt1-Cre, TrkA*^*Cre-ERT2*^, *Sst*^*Cre*^, *Vglut3*^*Cre*^, *Gfra3*^*CreERT2*^ and *Mrgprd*^*Cre*^ mice were crossed to *ROSA26*^*Tomato*^ and *ROSA26*^*ChR2*^ for histological characterization; and to *ROSA26*^*ChR2*^ for gain-of-function behavioral experiments. *TrkA*^*CreERT2*^ and *Gfra3*^*CreERT2*^ mice were crossed to *ROSA26*^*ArchT*^ for loss-of-function behavioral experiments. The resulting strains from crosses are as follows: *Wnt1Cre*;*ROSA26*^*ChR2*/+^ (Wnt1-ChR2), *TrkA*^*CreERT2*/+^;*ROSA26*^*ChR2*/+^ (TrkA^ChR2^), *TrkA*^*CreERT2*/+^;*ROSA26*^*ArchT*/*ArchT*^ (TrkA^ArchT^), *Mrgprd*^*Cre*/+^;*ROSA26*^*Tomato*/+^ (Mrgprd^TOM^), *Mrgprd*^*Cre*/+^;*ROSA26*^*ChR2*/+^ (Mrgprd^ChR2^), *Sst*^*Cre*/+^;*ROSA26*^*Tomato*/+^ (Sst^TOM^), *Sst*^*cre*/+^;*ROSA26*^*ChR2*/+^ (Sst^ChR2^), *Vglut3*^*Cre*/+^;*ROSA26*^*Tomato*/+^ (Vglut3^TOM^), *Vglut3*^*Cre*/+^;*ROSA26*^*ChR2*/+^ (Vglut3^ChR2^), *Gfra3*^*creERT2*/+^;*ROSA26*^*Tomato*/+^ (Gfra3^TOM^), *Gfra3*^*CreERT2*/+^;*ROSA26*^*ChR2*/*ChR2*^ (Gfra3^ChR2^), *Gfra3*^*CreERT2/*+^;ROSA26^ArchT/ArchT^ (Gfra3^ArchT^). All experiments were carried out in accordance with protocols approved by the Stockholm Ethical Committee for Animal Experiments (Stockholms Norra Djurförsöksetiska Nämnd, Sweden, 9702-2018 and 10406-2020). Animals were provided with food and water *ad libitum* and maintained on a 12-hour light/dark cycle.

For pups from mice crossed with TrkA^CreERT2^ and Gfra3^CreERT2^, Tamoxifen (Sigma, T5648) was dissolved in corn oil (Sigma, 8267) at a concentration of 20 mg/ml and delivered by intraperitoneal (i.p.) injection to P14 pups once and then in adult for two consecutive days (140 mg/kg both pups and adults). Control groups of test mice also received tamoxifen injections.

### Autoantibody-induced arthritis model

Arthritis was induced by intravenous (i.v.) injection of 6 mg cartilage autoantibody cocktail containing 4 arthritogenic monoclonal antibodies (ACC1: anti-citrullinated C1 epitope of collagen type II (COL2) antibody; M2139: COL2 antibody; L10D9: collagen type XI antibody; 15A: anti-cartilage oligomeric matrix protein antibody) on day 0 followed by 25 µg lipopolysaccharide (LPS, 055:B5, Sigma) intraperitoneally (i.p.) on day 5^31^. Control mice received 150 µL saline i.v. on day 0 while 100 µL saline i.p on day 5.

The mice were examined at different time points by arthritis scoring^98^. Briefly, each inflamed (both swollen and redness) digital was given score of 1 point and if dorsal side of the paw or wrist/ankle joint was inflamed 2.5 points were given for moderate inflammation and 5 points for severe inflammation, resulting in a maximum 15 for each limb and in total 60 per mouse.

### Light-induced response

A flexible optical fibre bundle monitored by power controller (DC2200, Thorlabs) was used to activate the channelrhodopsin2 (ChR2), and the withdrawal reflex was elicited using a pulsing laser (470 nm, 10 Hz, 50 ms ON/OFF) with intensities from low to high and applied onto the plantar surface of the hind paws. Wnt1-ChR2, TrkA^ChR2^, Sst^ChR2^, Vglut3^ChR2^, Gfra3^ChR2^ and Mrgprd^ChR2^ mice (n = 7-8) were habituated for 1 hour (h) on the mess floor, and a 20-second trial was conducted, alternating between the left and right hind paws with at least 10 minutes intervals.

For excitatory optogenetics, light threshold was determined as the lowest light power provoking a withdrawal response (for reflex) or nocifensive behaviour like shaking, lifting, licking and guarding (for coping) in one of stimulated hind paw. The percentage of withdrawal reflex responding mice in different strains was reported. In all experiments, subthreshold light stimulations (0.2% lower intensity than threshold) were applied simultaneously combined with the following described behavioral tests.

### Behavioral tests

For sensory behavioral tests, the mice were habituated to the test environment on two occasions before assessment of baseline. After two baseline recordings performed on different days, the animals were randomly assigned to saline control and arthritis groups. Mechanical sensitivity was determined by assessment of paw withdrawal using von Frey filament (Stoelting). A series of filaments with a logarithmically incremental stiffness of 0.04, 0.07, 0.16, 0.4, 0.6, 1.0 and 2.0 (g) was applied to the plantar surface of the hind paw and held for 3 sec according to the up–down method as previously described^99^. A brisk withdrawal of the paw was noted as a positive response. The 50% probability of withdrawal threshold (force of the von Frey hair to which an animal reacts to 50% of the presentations) was calculated as threshold. To avoid any potential tissue damage, a cut-off value of 2.0 g was applied. The average withdrawal threshold of two hind paws was used.

To assess heat sensitivity, a radiant heat source (IITC) was aimed at the plantar surface of the hind paw through a glass surface. Briefly, mice were placed in plexiglass cubicles on a glass surface. The thermal nociceptive stimulus originated from a projection bulb below the glass surface and the stimulus was delivered separately to one hind paw at a time. Latency was defined as the time required for the paw to show a brisk withdrawal. Each hind paw was tested three times with intervals and the average withdrawal latency was calculated.

Nocifensive episodes (paw shaking, lifting/guarding or licking) were measured as a quantitative scale of pain responses to a 2.0 g von Frey filament applied to both hind paws. Cold allodynia was assessed by calculating nocifensive episodes for 45 sec after application of one drop of acetone each to both hind paws. Mechanical hyperalgesia (pinprick) was tested with a safety pin (23G needle, BD), and Nocifensive behaviors were recorded. The average shaking numbers of two hind paws were used.

### Inverted screen test

To perform limb’s function, inverted screen test was measured: the mouse was placed in the center of the wire screen (width 7 mm and diameter 2 mm for wire, GMC500) and rotate the screen to an inverted position over 2 sec, with the mouse’s head declining first. Hold the screen steadily 45-50 cm above a soft material padded surface. Record the time mouse falls off (hanging time); animal will be removed from screen when it reaches the cut-off point (6 min).

### Clip squeeze test

For checking joint tenderness, after 1 hour of incubation in the Hargreaves’ box (IITC), a toothless clip (420 G) was applied to squeeze the proximal interphalangeal (PIP) joint and extension of the metatarsal-phalangeal (MTP) joint of one hind paw for 5 sec; then nocifensive episodes was analysed for 4 min after clip removal.

### Sunflower seed assay

To test the dexterity of forepaws, the sunflower seed assay was used, modified from previous descriptions^33, 100^. Animal was habituation to separated test box (animal enclosure, IITC) placed on the grey matte acrylic floor. After habituation, 2-3 sunflower seeds (provided by KM-B, KI) were applied to the floor for 20 min for 3 consecutive days (1 round of training). Only the activated mice (completely deshelling the seeds, around 60%) after 2 rounds of training were used for further study.

Two days prior to testing (habituation day 1 and day 2) and during testing day (testing on day 3), animals were transferred from their home cages and placed in the test box and allowed to explore the environment for 10 min. And then 2 seeds were placed on the floor and seed eating activity was recorded for 20 min. The episodes of rotation were calculated: the act of manipulating shell orientation rotating for 180 degrees within the forepaws.

For gain-of-function study, different pain-like behavior tests (mechanical and thermal) were applied before the measurement of light threshold; and then light threshold was determined as the lowest light intensity provoking a withdrawal (reflex) or nocifensive responses in one of the hind paws. Subthreshold light stimulations were then applied simultaneously with sensory stimuli: withdrawal reflex subthreshold for von Frey and Hargreaves tests and nocifensive subthreshold for 2g von Frey, acetone and pinprick tests. Note: the subthreshold light intensities for BL and arthritis mice were different, as they were adjusted for sensitization associated with arthritis (Wnt1-ChR2 mice BL reflex subthreshold: 12.7×10^−3^ mWatt/mm^2^, nocifensive subthreshold: 18.5×10^−3^ mWatt/mm^2^; Arthritis mice reflex subthreshold: 11.7×10^−3^ mWatt/mm^2^, nocifensive subthreshold: 14.5×10^−3^ mWatt/mm^2^; Mrgprd^ChR2^ mice no difference in subthreshold light; TrkA^ChR2^ mice BL reflex subthreshold: 19.7×10^−3^ mWatt/mm^2^, Arthritis mice reflex subthreshold: 15.4×10^−3^ mWatt/mm^2^). No effects were found in Sst^ChR2^ or Vglut3^ChR2^ mice with the subthreshold gain-of-function blue light in normal or arthritis mice.

In littermate control mice (Wnt1-Cre, TrkA^CreERT2^ or ROSA26^ChR2^ mice), optogenetic activation did not induce stimuli-associated responses, neither withdrawal nor shaking behavior. All three strains showed mechanical and cold hypersensitivity, however no differences of sensitivities to mechanical or thermal stimuli were found in these control mice comparing with and without light stimulation (data not shown).

### Inhibitory optogenetics

To test the effect of inhibition of TrkA and *Gfra3* expressing sensory neurons, mechanical and thermal sensitivities were assessed as well as joint pain tests (clip squeeze and inverted screen tests and sunflower seed assay) before and after yellow light (30 min for TrkA^ArchT^ and 45 min for Gfra3^ArchT^ mice, n = 8) stimulation. A lab-made yellow LED-plate (wavelength: 566 nm; 0.1 mWatt/mm^2^) was positioned under the testing floor. For mechanical sensitivity, after 1-hour habituation on a mesh floor, the plantar surface of the hind paws was stimulated with a series of calibrated monofilaments (Stoelting) with increasing force (0.07 g, 0.16 g, 0.4 g, 0.6 g, 1.0 g, 1.4 g and 2.0 g). Each filament was applied five times to both hind paws. The percentage of animals with withdrawal reaction was reported.

### DRG single cell suspension preparation

C57BL/6N mice (8-10 wk, Charles River, Sweden) were sacrificed with overdosage of isoflurane. Spines were removed and fresh cervical and lumbar dorsal root ganglia (DRGs) were dissected in a 6 cm Petri-dish with DPBS (Sigma) on ice. And 2 male mice were included for each suspension experiment. The following DRG single cell suspension was performed according to our previous protocol with modifications^101^. Briefly, dissected DRGs were chopped with microscissors 1-2 times in 2 mL Papain (25 unit/mL, Worthington Biochemical) and then incubated at 37 °C, with a mixture of digestion enzymes of Papain/Collagenase/Dispase (Papain, 25 unit/mL, 4 mL; DNase I, 55 unit/mL, 0.5 mL, Worthington Biochemical; and Collagenase and Dispase 20 mg/mL, 800 µL, Worthington Biochemical). The DRGs were triturated up and down 10 times of every 10 min using glass Pasteur pipettes with decreasing diameter (pre-coated with 0.5% BSA). Cell suspension was filtered through a 30 µm cell strainer (CellTrics, Sysmex) and washed with additional 1.5 mL artificial cerebrospinal fluid (ACSF, 87 mM NaCl, 2.5 mM KCl, NaH_2_PO_4_ 1.25 mM, NaHCO_3_ 26 mM, Sucrose 75 mM, Glucose 20 mM, CaCl_2_ 0.5 mM, MgSO_4_ 4 mM) and 0.5 mL DPBS. Cells were pelleted by centrifugation (300 g × 6 min, 4 °C) and resuspended with 1.5 mL cold ACSF with 0.5 mL DPBS. The cell suspension was carefully loaded on top of the same volume of OptiPrep density gradient medium (Sigma) and centrifuged with 700 g ×10 min at 4 °C. The cell pellet was resuspended with 3 mL cold ACSF. SYTOX Blue (Invitrogen, ThermoFisher Scientific) was added to stain dead cells. Thereafter, alive SYTOX Blue-negative cells were sorted with fluorescence activated cell sorting (FACS, BD FACSAria Fusion / BD FACSAria III) at 4 °C. Cells were concentrated by centrifugation (300 g × 5 min, 4 °C) and resuspended with proper volume (∼1 000 cells/ µL) of ACSF solution.

### Single cell gene expression 3’ sequencing

Sorted cells were loaded onto the 10× Chromium chip G to yield single cell droplet with v3.1 kit (10× genomics). Targeted cell recovery for each well was settled to 5 000 cells (∼10% neurons). Reverse transcription, cDNA amplification and library construction were performed according to the user guide provided by 10× genomics. Pooled libraries were sequenced on Illumina sequencing platform NovaSeq 6000 system on SP-100 flowcells with 91 bp sequenced into the 3’ end (5’ to 3’) of the mRNAs in National Genomics Infrastructure (SciLifeLab). Raw sequencing data were de-multiplexed, converted into fastq format, and aligned to mouse reference mm10 using the STAR aligner to generate the gene-cell matrices.

### sc-RNA sequencing data analysis

R (v.4.1.1) using Seurat (v.4.1.0) was used for the main scRNA-seq analysis. Individual count matrices created by CellRanger (v.5.0.1) were merged to a single Seurat object and all cells with more than 20% of counts originating from mitochondrial genes were discarded. A cutoff at more than 2 000 detected genes was set for the primary data. These data were integrated using Harmony (v.0.1.0) and clustered using the default algorithm in Seurat. Putative neuronal clusters were identified using the neuronal marker gene *Rbfox3*. Nonneuronal clusters from control samples were extracted, integrated and clustered, and assigned cell labels based on gene markers from literature ^102^. The nonneuronal control data was then used to transfer labels (Seurat) to all remaining nonneuronal data. After this, all remaining original data with > 1 000 detected genes were integrated, clustered and assigned labels from the primary data. All neurons from this secondary data were discarded to make sure that only high-quality neurons were used for the final analyses. Then, the primary (> 2 000 detected genes) and secondary ≥ 1 000 detected genes) datasets were merged to produce the full working dataset. More granular identities for the immune cells in the data were assigned using a peripheral nerve immune cell atlas^102^. For this, a mixture discriminant analysis (mda) based classifier (scPred, v.1.9.2) was built using these data and the cell type labels for the immune cells in the present data were learned using this model. All cells with a prediction score below 0.55 were discarded. For neurons, all cells labeled as neurons were extracted from the full working data, clustered and using iterative clustering steps removing all cells with less than 0.5 normalized counts of *Rbfox3* and more than 2 normalized counts of *Apoe*. A classifier was then built as before, using data from Zeisel *et al*. with Usoskin *et al*. annotation and the cell type labels for the neuronal data were learned using the model and unassigned neurons discarded similarly as stated above. For a pseudobulk differential expression (DE) analysis, the neuron types were collapsed together and data from each individual timepoint after inducing arthritis were compared against control (t0) using Wilcoxon Rank-Sum test with the Seurat function FindMarkers with adj.p.val cutoff set at 1×10^−20^. DE genes for each cell type between individual arthritis timepoint and control were defined in a similar fashion. Fcoex (v.1.10.0) was used to identify co-regulated gene modules in the dataset. For this, to reduce computational load, a random set (25%) of cells from each cell type-timepoint pair was sampled. Fcoex was run for the first 200 genes using “timepoint” as the target. The resulting set of modules was further filtered to contain only differentially expressed genes and modules that consisted of minimum 10 genes. A module score was calculated for modules and scaled to fall between 0 and 1. A gene enrichment analysis for gene modules was run using enrichR (v.3.0) with the “GO_Biological_Process_2021” and “KEGG_2019_Mouse” databases. For the perturbation analysis (Augur v.1.0.0), all genes situated on the Y-chromosome and non-protein-coding genes were first discarded. Following this, the analysis was run comparing each individual timepoint against control for each neuron type. The default minimum of 20 cells per type/timepoint was used; therefore, some neuron types were not compared for each timepoint.

### Intercellular ligand-receptor analysis

We identified upregulated genes at the 12h timepoint compared to the 0-hour timepoint by applying the following criteria: a fold change of at least 3 times, gene expression in at least 10% of cells within a cell type, and a p-value of less than 0.05. The upregulated ligand genes were further selected from these increased genes, and all expressed genes were used for determining the receptor genes in each cell-type.

We next calculated the ligand-receptor pairs according to the ligand-receptor dataset as used^103^. We then collected associated gene patterns for each ligand-receptor pair from nichenetr and SCENIC^53, 104^. We evaluated the ligand-receptor activity by scoring the associated gene patterns of each ligand-receptor pair upregulated at the 12h timepoint using the enrichment score^105^. Briefly, we used all upregulated genes as the candidate gene pool. Ligand-receptor associated gene patterns were selected for scoring, while the remaining genes were considered background genes and ranked based on their expression. We created 20 intervals according to the background gene expression and randomly selected 100 genes from each interval to form a random background gene matrix. Feature scores were calculated by comparing the average expression of the associated genes to the randomized background genes. All negative values were normalized to 0, indicating non-activity. To reduce variation, for each ligand-receptor pair between two cell types, we repeated the enrichment score calculation five times and determined the average value as the activity score. The dot map was visualized to reflect the ligand-receptor activity of all ligand-receptor pairs from lig- and-expressing cells (sender cell types) to receptor-expressing cells (receiver cell types), with colours ranging from blue to red indicating activity from low to high. The specific ligand-receptor pairs between different cell types were visualized in a Sankey plot. The codes and analysis steps will be accessible as a Jupyter Notebook on https://sccamel.readthedocs.io/.

### Type I IFN signaling blocking

C57BL/6N mice received either a neutralizing monoclonal antibody against IFNAR1 (40 mg/kg, i.p., BioXCell) or an isotype of mouse IgG1 antibody (40 mg/kg, i.p., BioXCell) 1 hour prior to the injection of cartilage autoantibodies (day 0), or on day 22, or on day 45 after autoantibody injection. Mechanical sensitivity was tested 12h (0.5d), 36h (1.5d), 60h (2.5d) and 84h (3.5d) after IFNAR1 mAb or control isotype mAb injection (n = 6). For joint function study, IFNAR1 or isotype (40 mg/kg, i.p.) was injected 1 hour before autoantibody cocktail administration, or on day 11, or day 33 after antibody injection. And inverted screen test, sunflower seed assay and squeeze test were measured 12h (0.5d), 36h (1.5d), 60h (2.5d) and 84h (3.5d) after IFNAR1/Isotype treatment (n = 5). TYK2 inhibitor (Deucravacitinib/MBS-986165, MCE) was orally administrated for 9 days from day 13 after autoantibody injection in C57BL/6N mice twice daily (8 AM and 8 PM, 15 mg/kg, in EtOH:TPGS:PEG300 of 5:5:90)^106^. Mechanical sensitivity of von Frey withdrawal threshold and 2g von Frey nocifensive behavior was measured 1.5h after morning injection of TYK2 inhibitor or vehicle (EtOH:TPGS:PEG300 for 5:5:90) (n = 6). A single i.p. injection of MNK1/2 inhibitor, Tomivosertib (eFT508/HY-100022, MCE) on day 48 after antibody injection in C57BL/6N mice (1 mg/kg, in DMSO:PEG300:Tween-80:Saline of 5:40:5:50), and then mechanical sensitivity of von Frey withdrawal threshold as well as nocifensive behavior of 2g von Frey and clip squeeze tests were measured around 1.5h and 24h after Tomivosertib administration; and on day 55 sunflower seed assay and inverted screen test were checked around 1.5h and 24h after Tomivosertib administration (n = 10, 5 females and 5 males). eIF4E/eIF4G interaction inhibitor, 4EGI-1 (324517, Sigma) was i.p. injected into antibody-induced arthritis C57BL/6N mice (15 mg/kg, in DMSO:PEG300:Tween-80:Saline of 5:40:5:50, on day 56). Mechanical sensitivity (von Frey up-down and 2g von Frey nocifensive tests) as well as clip squeeze test, sunflower seed assay and inverted screen test were checked (n = 5).

### Endo S treatment of autoantibody

For the Fc N-glycan cleavage, GST-fused endoglycosidase S (EndoS) expressed by *E*.*coli* was used to incubate with the cartilage antibody cocktail at a ratio of 1:1 000 (w/w) and 37 °C for 1h. All antibodies were purified by using Protein G GraviTrap Columns (Cytiva) according to the manufacturer’s instructions.

### Cytokine measurements in serum

Mice were deeply anesthetized with sodium pentobarbital (60 mg/kg) and blood was collected via heart. After sitting at room temperature for 30 min, blood samples were centrifuged 1 000 g × 10 min at 4 °C, and serum samples were aliquoted from supernatant and stored at -80 °C until use. Serum levels of interferon alpha (IFNα) and interferon beta (IFNβ) as well as other 31 cytokines/chemokines were measured by multiplex assay service (Eve technologies) using Mouse IFN 2-Plex Discovery Assay (MDIFNAB) and Mouse Cytokine/Chemokine 31-plex Discovery Assay Array (MD-31), respectively (n = 4).

### Quantitative PCR of interferons in DRG samples

Fresh DRG samples were collected at different timepoints after autoantibody injection: 1h, 12h, 3d, 33d and 63d, 4 mice for each group. Naïve C57BL/6N mice (n = 9) were used as the control group. For IFNAR1 treatment group, samples were harvested on 33d after 12h treatment of IFNAR1 mAb (40 mg/kg, i.p., BioXCell) (n = 5). Fresh frozen human lumbar DRGs were provided by AnaBios (detailed information in supplementary Table S8). Total RNA was extracted from mouse cervical and lumbar DRGs using TRIzol Reagent (Thermo Fisher Scientific) and Motorized Pestle Mixer (Argos Technologies) as previously described^107^. cDNA was generated from 500 ng RNA using High-Capacity cDNA Reverse Transcription Kit (Applied Biosystems) with random primers according to the manufacturer’s instructions. qPCR reactions were performed using SYBR Green Master Mix (Thermo Fisher Scientific) on a QuantStudio5 System (Applied Biosystems). Primer pairs used in this study were the following: for mouse genes, *Ifna* (pan primers, forward: CCTGAGAA/GAGAAGAAACACAGCC, reverse: GGCTCTCCAGAC/TTTCTGCTCTG), *Ifnb* (forward: AGGGCG-GACTTCAAGATC, reverse: CTCATTCCACCCAGTGCT), *Il1b* (forward: GAATCTATACCTGTCCTGTG; reverse: TTATGTCCTGACCACTGTTG), *Il6* (forward: CTCATTCTGCTCTGGAGCCC; reverse: TGCCATT-GCACAACTCTTTTCT), *Tnfa* (forward: GACCCTCACAC-TCAGATCATCT; reverse: CCTCCACTTGGTGGTTTGCT), *Gapdh* (forward: AACTTTGGCATTGTGGAAGG; reverse: ACACATTGGGGGTAG-GAACA); for human genes, pan-IFNA (forward: TCCATGA-GATGATCCAGCAGA; reverse: ATTTCTGCTCTGACAACCTCCC), IFNB (forward: CAACTTGCTTGGATTCCTACAAAG; reverse: TATTCAA-GCCTCCCATTCAATTG), IL1B (forward: TTCTTCGACACATGGGA-TAACG; reverse: TGGAGAACACCACTTGTTGCT), IL6 (forward: TAATGGGCATTCCTTCTTCT; reverse: TGTCCTAACGCTCATACTTTT), TNF (forward: CAGCCTCTTCTCCTTCCTGAT; GCCAGAGGGCTGATTAGAGA), GAPDH reverse: (forward: GCCAAGGTCATCCATGACAACTTTGG; reverse: GCCTGCTTCACCAC-CTTCTTGATGTC). All assays were performed in duplicate, and the levels of transcripts were analyzed by the comparative CT (2^-Δ ΔCt^) method relative to *Gapdh/*GAPDH.

### Tissue preparation and in situ hybridization (RNAscope)

Mice were deeply anesthetized with isoflurane and decapitated. Lumbar DRGs was dissected out, snap frozen on dry ice and kept at −80 °C. Fresh frozen human lumbar DRGs were provided by AnaBios (L3-L5 DRGs, Supplementary Table 8). Frozen tissues were thawed on ice briefly, trimmed, embedded with optimal cutting temperature cryomount, frozen with liquid carbon dioxide, and sectioned on a cryostat at a thickness of 12 µm for DRGs. The sections were mounted onto Superfrost Plus microscope slides and stored at −80 °C until use.

RNAscope assay was performed according to the protocol provided with RNAscope Multiplex Fluorescent Detection Kit v2 (Cat. No. 323110, ACDBio) with minor modifications, where the hydrogen peroxide treatment was skipped, protease III was used instead of protease IV and counterstaining with DAPI (1.0 µg/ml, 10 min at RT) was performed. Slides were mounted after rinse in PBS with mounting medium (Agilent Dako), dried overnight at RT and stored at −20 °C until imaging. The probes included in this study were designed and provided by ACDBio as listed here (Probe-Mm-*Ifnas*-cons-o1 cat# 471691, Probe-Mm-*Ifnar1*-C2 cat# 512971, Probe-Mm-*Calca*-alltv-C3 cat# 417961, Probe-Hs-*TRPV1*-C2 cat# 415381, Probe-Hs-*IFNAR1*-C3 cat# 500891, Probe-Hs-*IFNAR2* cat# 490151, Probe-Hs-*SOX10*-C2 cat# 484121).

### Immunohistochemistry

Mice were deeply anesthetized with sodium pentobarbital (60 mg/kg) and perfused transcardially with 20 mL pre-warmed (37 °C) saline, followed by 20 mL of pre-warmed 4% paraformaldehyde in 0.16 M phosphate buffer (pH 7.2-7.4) and 50 mL cold fixative. L4/L5 DRGs were dissected and post-fixed in the same fixative for 90 min at 4 °C. After cryoprotection in 10% sucrose with 0.1 M phosphate buffer containing 0.01% sodium azide (VWR) and 0.02% bacitracin (Sigma) for 48h, the tissue was embedded with OCT (His-toLab), frozen with liquid carbon dioxide and sectioned on a CryoStar NX70 cryostat (Thermo Scientific) at 12 µm thickness.

For staining, mounted sections were dried at room temperature for at least 30 min and then incubated with antibodies against NF200 (1:1 000, chicken, Abcam), CGRP (1:10 000, rabbit, gift from Tomas Hökfelt), GFP (1:1 000, chicken, Abcam) and TH (1:200, rabbit, Pel-Freez) diluted in phos-phate-buffered saline (PBS) containing 0.2% (wt/vol) BSA (Sigma) and 0.3% Triton X-100 (Sigma) in a humid chamber at 4 °C for 48h. Immunoreactivities was visualized with secondary IgG (H+L) antibody conjugated with Alexa Fluor™ dyes (Invitrogen) at room temperature for 90 min. For IB4 staining, slides were rinsed in PBS for 20 min and incubated with IB4 (1:400) from Griffonia simplicifolia I (GSA I) (2.5 g/mL; Vector Laboratories), followed by overnight incubation with a goat anti-GSA I antiserum (1:2 000; Vector Laboratories). Finally, the sections were incubated with IgG (H +L) secondary antibody conjugated with Alexa Fluor (Invitrogen) to visualize the IB4 binding. DRG Sections from Gfra3^TOM^ mice were counterstained with DAPI (Sigma) for 10 min at room temperature. After rinsing in PBS, the sections were mounted with fluorescence mounting medium (Agilent Dako).

### Microscopy and image processing

Representative images were acquired from one airy unit pinhole on an LSM700 confocal laser-scanning microscope (Carl Zeiss) equipped with EC Plan-Neofluar objectives with magnifications of 10× and 20×. Emission spectra for each dye were limited as follows: DAPI (< 480 nm), Alexa Fluor 488 (505–540 nm), Alexa Fluor 555/PI/Cyanine 3 (560–610 nm), and Alexa Fluor 647/Cyanine 5 (> 640 nm). Images were processed using ZEN software (Zeiss). Multi-panel figures were assembled using Adobe Photoshop 2022 and Adobe Illustrator 2022 (Adobe Systems).

### Patch-clamp electrophysiology

Cell cultures for patch-clamp electrophysiology were prepared from adult C57BL/6N mice (both sexes, 7-10 weeks). Briefly, all levels of cervical and lumbar DRGs were dissected and digested in Papain/Collagenase/Dispase mixture, then triturated using glass Pasteur pipettes. Dissociated cells were suspended in L-15 medium (Liebovitz, L1518, Merck) contained 10% FBS, NaHCO_3_, Glucose, penicillin/streptomycin (1×) and Floxuridine (PHR2589, Merck) and plated on poly-D-lysine (A-003-E, Merck) and laminin (L2020, Merck) pre-coated coverslip. The following day, changes in neuronal excitability in small-sized nociceptors, (diameter ≤20 µm) were tested after 1 hour preincubation in recombinant mouse IFNa3 protein (12100-1, 300 U/mL, B&D systems), PBS containing 0.1% BSA (Sigma) stimulation in neurons served as controls. To test the effects of MNK1/2 inhibition on IFN-stimulated nociceptors, neurons were first incubated with 10 µM eFT508 (in L-15 medium) or vehicle (DMSO, 0.1% v/v) for 1 hour and IFNα3 (300 U/mL) was then added. After 1 hour IFNα3 incubation, the coverslip was placed in a 35 mm petri dish filled with Artificial cerebrospinal fluid (ACSF) solution. ACSF was composed of 125 mM NaCl, 25 mM glucose, 25 mM NaHCO_3_, 2.5 mM KCl, 2 mM CaCl_2_, 1.25 mM NaH_2_PO_4_, 1 mM MgCl_2_ that was saturated with 95% oxygen and 5% carbon dioxide and maintained at room temperature (20-22 °C).

All recordings were performed using whole-cell patch-clamp technique. Patch-clamp electrodes were filled with a solution containing (in mM): 120 K-gluconate, 5 KCl, 10 HEPES, 4 Mg_2_ATP, 0.3 Na_4_GTP, 10 Na-phosphocreatine with pH 7.4 adjusted with KOH and an osmolarity of 275 mOsm. Neurons were visualized using a fluorescence microscope (Axioskop FS Plus, Zeiss) equipped with IR-differential interference contrast (DIC) optics and a CCD camera (Hamamatsu). Patch-clamp electrodes were advanced into the dish using a motorized micromanipulator (Luigs & Neumann) while applying constant positive pressure. Intracellular signals were amplified using a MultiClamp 700B amplifier (Molecular Devices) and low-pass filtered at 10 kHz. Electrophysiological data was digitized at 10 or 20 kHz using a Digidata 1322A A/D converter (Molecular Devices) and acquired using pClamp software (Molecular Devices). Neurons were held at -60 mV in current-clamp mode and ramps of depolarizing current (1s duration, peak amplitudes of 100, 300, 500 and 700 pA) were injected into each neuron. The total number of action potentials elicited by each current ramp was quantified for each neuron.

### Western blotting

To perform western blot in arthritic tissues, cervical and lumbar DRGs were collected at different time points after antibody injection: 2h, 12h, and 33d,4 mice for each group. For IFNAR1 treatment group, samples were harvested on 33d after 12h treatment of IFNAR1 mAb (40 mg/kg, i.p., BioXCell) (n = 3). Naïve C57BL/6N mice (n = 9) were used as the control group. Total protein was extracted from mouse DRG samples using N-PER neuronal protein extraction reagent (87792, ThermoFisher) containing Protease inhibitor cocktail (G6251, Promega) and Halt Phosphatase inhibitor cocktail (78428, ThermoFisher). Tissues were homogenized using micropestle (Sigma) and followed by sonication and then sit on ice for 10 min. Centrifuge at 10,000 g for 10 min at 4°C. Supernatants were collected and the total protein concentration was measured by BCA assay kit (ThermoFisher). Protein lysates from fresh human lumbar DRGs from healthy donors, painful RA patients and RA patients without pain donors (AnaBios, Supplementary Table 8) were homogenized with bead using Tissue homogenizer (Qiagen) and then processed in a similar way as mouse DRGs. For mouse DRG lysates, 20 µg of denatured protein was separated by electrophoresis on a NuPAGE 4-12% Bis-Tris gel and transferred onto 0.2 µm nitrocellulose membrane by iBlot2 transfer system. Membranes were blocked with 5% BSA in TBST (0.1% Tween 20) for 1hr at RT and then incubated with primary antibody against Phospho-eIF4E (Ser209, 1:1,000; #9741, CellSignaling technology) for two days at 4°C. Membranes were washed in TBST for 3’ 15 min at RT on a shaker and incubated with polyclonal secondary antibody conjugated with horseradish peroxidase (HRP) in 5% BSA (1:10,000; P039901, DAKO) for 1 hr at RT. The signal was detected with SuperSignal West Femto reagents (1:1 diluted with water, ThermoFisher) after washing the membrane and imaged with a ChemiDoc MP system (Bio-Rad Laboatories). Membranes were stripped in Restore Plus Western Blot stripping buffer (#46430, ThermoFisher) for 1 hr, washed with TBST (3 × 15 min), blocked with 5% BSA and re-probed with primary antibody against eIF4E (1:1,000; #9742, CellSignaling technology) overnight at 4°C. The membrane was washed, incubated with HRP conjugated secondary antibody, detected with Amersham ECL prime Western Blotting detection reagents and imaged by the ChemiDoc system. GAPDH (1:5,000; #5174, CellSignaling technology) was probed on the stripped membranes as the loading control in the end. Band intensity of specific protein bands were quantified with Image Lab 6.1 software (Bio-Rad laboratories). Protein phosphorylation level was normalized to total protein expression and compared to normalized control samples. For human DRG lysates, 30 µg of denatured protein was loaded for western blotting and probed with primary antibodies (IFN alpha from ThermoFisher, PA5-86767; beta-Actin from Abcam, ab6276) in a similar protocol as described above.

### Ex vivo teased tibial nerve recordings

Extracellular recordings from single cutaneous primary afferent axons in an isolated mouse glabrous skin–tibial nerve preparation was obtained following previously published procedures^108, 109^. In brief, around 3 months (day 85-day 98) after autoantibody (Arthritis) or saline (Control) injected mice (n = 5, both males and females) were euthanized by cervical dislocation and the glabrous skin from one hind paw with the tibial nerve attached was dissected and placed in a custom made two-compartment teflon recording chamber with the corium side down. The chamber containing the preparation was continuously superfused at a rate of 5 mL/min with oxygenated external solution consisting of: 107.8 mM NaCl, 26.2 mM NaHCO_3_, 9.64 mM sodium gluconate, 7.6 mM sucrose, 5.55 mM glucose, 3.5 mM KCl, 1.67 mM NaH_2_PO_4_, 1.53 mM CaCl_2_ and 0.69 mM MgSO_4_, which was adjusted to pH 7.4 by continuously gassing with 95% O_2_–5% CO_2_. Temperature ± 1 °C using a heat exchanger connected to a thermostat^110^. Tibial nerve was placed into an adjacent chamber of the bath filled with mineral oil and then teased into small bundles that were individually placed on a gold wire electrode. A reference electrode was positioned inside the recording chamber dipped into the aqueous solution. Input signals were amplified through a high gain AC differential amplifier (Neurolog NL104A; Digitimer), digitized (PowerLab 8; ADInstruments) at 25 kHz and stored in the hard drive of a PC for offline analysis. LabChart software package (ADInstruments) was used for recording and off-line analysis. Mechanically responsive receptive fields were identified by probing the skin flap with a blunt glass rod. Once a suitable fiber was found a mechanical stimulator consisting of a tension/length feedback controller (300C-I; Aurora Scientific) was used to apply mechanical stimuli. Two different force protocols were used to characterize mechanical responses. Threshold and firing frequencies were measured during continuous force ramps from 0 to 100 mN (ramp duration 10 sec). Firing frequencies were also recorded during static force applications from 0 to 5, 10, 20, 40, 50, 75, 150, and 200 mN (step duration 10 sec; 50 sec interforce interval). Only mechanically responsive C fibers (conduction velocity < 1.2 m/s) were used in these experiments^111^. Experimenter was blinded to genotype until data analysis was complete.

### Statistics

Clinical score was shown as mean ± standard error of mean (SEM), and behavior data for von Frey filament up-down test and nocifensive behavioral tests including 2g von Frey test, acetone cold allodynia, pinprick and squeeze tests as well as sunflower seed assay (non-continuous data) were presented as median with interquartile range. Heat hypersensitivity behavioral data was presented as mean with SEM. Inverted screen test data, western blotting quantification data and cytokine levels in sera were presented as mean ± standard deviation (SD). qPCR data was shown as mean ± SEM. A *p* value less than 0.05 was considered significant (* indicates *p* < 0.05; ** indicates *p* < 0.01 and *** indicates *p* < 0.001). Data was analyzed with Prism 10.2.0 (GraphPad software) as specified in figure legends.

## Acknowledgements

We thank Cheryl Stucky (University of Minnesota) for generously sharing design and troubleshooting of optogenetic setups used in this study. We thank animal staff (Heidi Rautiainen, Tamara Teplova, Felicia Köhler, Isabo Gustafsson, Marina Sarvanidou and Ulrica Åberg) from Department of Comparative Medicine helping with animals (KM-B, Karolinska Institutet), laboratory managers Jana Sontheimer and Roland Baumgartner for animal genotyping and technical supports, Biomedicum Flow Cytometry Core facility (Karolinska Institutet) supported by KI/SLL for providing cell sorting services. We thank Prof. Jorge Ruas (Karolinska Institutet) for providing Tissue-homogenizer. We acknowledge support from Biomedicum Imaging Core (BIC) facility at Karolinska Institutet, supported by the Knut and Alice Wallenberg Foundation. We acknowledge the Eukaryotic Single Cell Genomics (ESCG) Facility at the Science for Life Laboratory, Sweden, for the scRNA-seq. The computations and data handling were enabled by resources in projects SNIC 2022/23-445 and -848 provided by the National Academic Infrastructure for Supercomputing in Sweden (NAISS) at UPPMAX, funded by the Swedish Research Council through grant agreement no. 2022-06725.

This work was supported by the Swedish Medical Research Council (2019-00761), Knut and Alice Wallenbergs Foundation (Wallenberg Scholar and Wallenberg project grant, KAW 2016-0006, 2019-0275), Wellcome Trust (Pain Consortium), European Research Council advanced grant (DescendPain 101053091), and Karolinska Institutet (to P.E.); the Swedish Medical Research Council (2019-01209), Knut and Alice Wallenbergs Foundation (2019-0059) (to R.H.); Åke Wibergs Grant to Y.H.; Brain foundation postdoctoral fellowships (to J.S. and M-D.Z.) and Swedish Society for Medical Research (SSMF) postdoctoral fellowships (to J.S. and M-D.Z.).

## Author contributions

J.S., M-D.Z., J.K., R.H. and P.E. conceived experiments. J.S., M-D.Z., J.K. and P.E. designed experiments. J.S., M-D.Z., J.K., A.E.M, R.H. and P.E. interpreted the results and wrote the paper. J.S. did the mouse genetics, all animal work, behavioral analyses and qPCR experiments. M-D.Z. collected tissue, generated scRNA-seq libraries, run western blot in mouse DRG tissue and did IHC and RNAscope experiments. J.K. performed the computational analyses of scRNA-seq datasets. D.K. run western blot in human DRG tissue. L.P. did patch-clamp in mouse DRG neurons. B.Z.X. purified autoantibodies. A.G.A performed and analyzed the single nerve recordings. Y.H. analyzed ligand-receptor intaeraction. D.U. built the equipment for optogenetic excitation and inhibition. Z.W.X. generated and purified EndoS autoantibodies. J.S., M-D.Z., J.K., R.H. and P.E. wrote the paper with input from all authors.

## Additional information

Supplementary information ExtendedData Figures 1 - 10, Supplementary Figures 1 - 3, Supplemental tables 1 - 8, Supplemental videos 1 - 2.

