## Supplementary material for "Persistent interferon signaling that causes sensory neuron plasticity and pain in arthritis": Su et al_2025_Supporting Information.pdf

bioRχiv

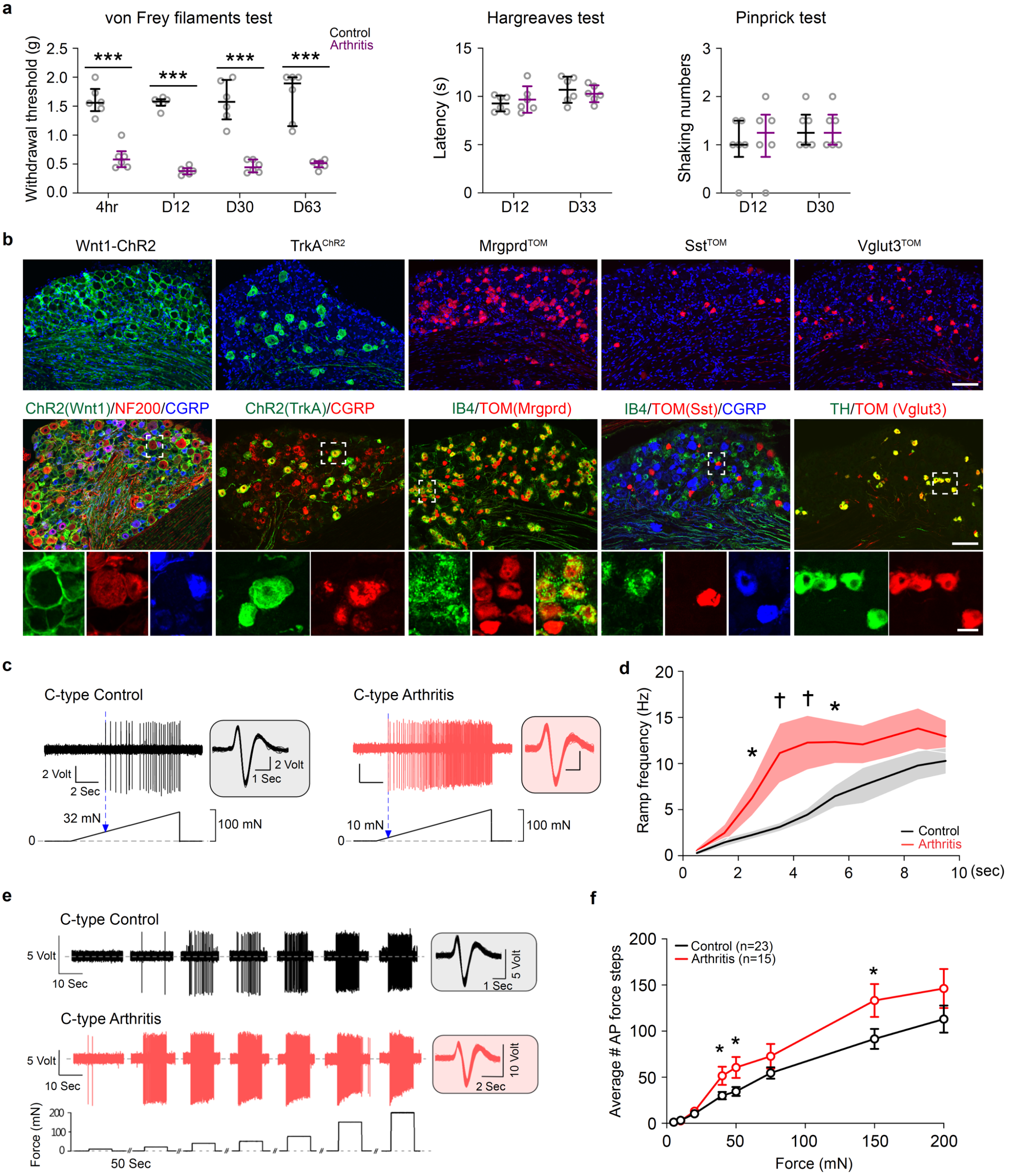


**Extended Data Fig. 1 Mechanical hypersensitivity in cartilage autoantibody-induced arthritis mouse.** (**a**) Mechanical allodynia in cartilage autoantibody-induced arthritis mice model with C57BL/6N mice from acute to chronic phases tested by von Frey up-down test. Heat sensitivity was tested by Hargreaves test and mechanical hyperalgesia was tested by pinprick test (n = 6 in each group, ***$p$<0.001). Kruskal-Wallis test followed by Dunn's multiple comparisons test for von Frey filaments and pinprick tests, one-way ANOVA followed by Dunnett multiple comparisons test for Hargreaves test. (**b**) Characterization of mouse strains Wnt1-ChR2, TrkA$^{ChR2}$, Mrgprd$^{TOM}$, Sst$^{TOM}$ and Vglut3$^{TOM}$ with immunohistochemistry comparing reporter expression (Tom) and markers for neuronal subtypes as indicated. NF200, Neurofilament 200; CGRP, Calcitonin-gene related peptide; IB4,

Isolectin B4; TH, Tyrosine hydroxylase. Scale bars = 100 μm and 20 μm for the inset. (**c**) Representative C-type fiber recordings (CV < 1.2 m/s) showing the activity during force ramp applications (10 sec; 0 to 100 mN) from mice injected with saline ("C-type Control"; left panel) or cartilage autoantibody ("C-type Arthritis", 3 months after antibody injection; right panel), the insets show all the action potential waveforms elicited during the mechanical stimulus. Bottom: force ramp protocol used to determine the mechanical threshold in individual fiber. (**d**) Mechanically induced firing frequencies of from control and arthritis mice during the force ramp application, thick lines represent mean number of action potentials in 1 second bins, shadowed regions are the SEM (unpaired *t* test, *$p<0.05$, †$p<0.005$). (**e**) Representative C-type fiber recordings during static force application from control and arthritis mice, the insets show all the action potential waveforms elicit during the mechanical stimulus. (**f**) Average number of mechanically induced action potentials during the static indentation (unpaired *t* test, *$p<0.05$).

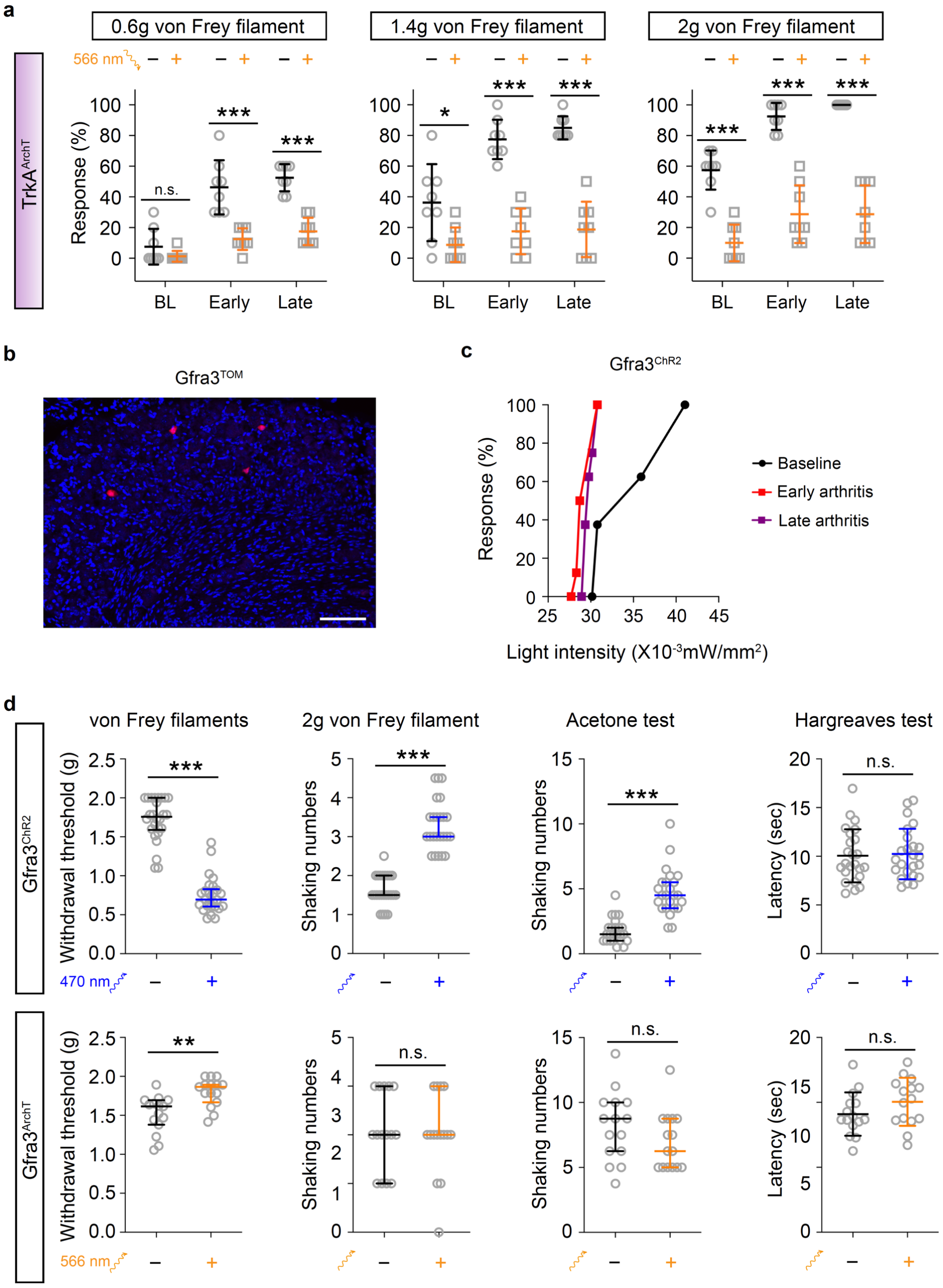


**Extended Data Fig. 2 The role of *Gfra3*+ C-fiber nociceptors in normal mechanical and cold sensation.** (**a**) In naïve *$TrkA^{CreERT2}$;$ROSA26^{ArchT}$* ($TrkA^{ArchT}$) mice, exposure to yellow light (0.1 mWatt/mm$^2$, 30min) caused less percentage of reflex reaction to 1.4 g and 2.0 g von Frey filaments indicating that TrkA neurons contributes to noxious, but not innocuous (0.6 g) pricking. In autoantibody-induced arthritis mice, inhibition of the TrkA population of neurons by stimulation of yellow light (0.1 mWatt/mm$^2$, 30min) completely reversed the increased sensitivity and reflex reactions in both early and late phases of arthritis (n = 8, unpaired *t* test, **p*<0.05, ****p*<0.001). (**b**) *Gfra3*+ neurons (Tom) in DRGs of *$Gfra3^{CreERT2}$;$ROSA26^{Tomato}$* ($Gfra3^{TOM}$) mice, DAPI (blue) is nuclear counter staining. Scale bar = 100 μm. (**c**) Percentage of reflex responses in mice to blue light at different light intensities in *$Gfra3^{CreERT2}$;$ROSA26^{ChR2}$* ($Gfra3^{ChR2}$) mice, with left-shifted response curve in early and late phases of antibody-induced arthritis (n = 8). (**d**) In $Gfra3^{ChR2}$ mice, a combination of blue light (subthreshold for reflex: 30.2×10$^{-3}$ mWatt/mm$^2$; subthreshold for nocifensive: 39.6×10$^{-3}$ mWatt/mm$^2$) increased mechanical and cold sensitivity with lower mechanical threshold and more nocifensive episodes to 2g von Frey filament and acetone test but had no effect on thermal sensitivity (latency in Hargreaves) (n = 25, ****p*<0.001). In *$Gfra3^{CreERT2}$;$ROSA26^{ArchT}$* ($Gfra3^{ArchT}$) mice inhibition by yellow light (0.1 mWatt/mm$^2$, 45 min) led to higher mechanical withdrawal threshold in von Frey test compared to before exposure to yellow light (n = 16, ***p*<0.01). Mann-Whitney test was applied for von Frey filaments, 2g von Frey filament and acetone tests, whereas Hargreaves test was analyzed by unpaired *t* test.

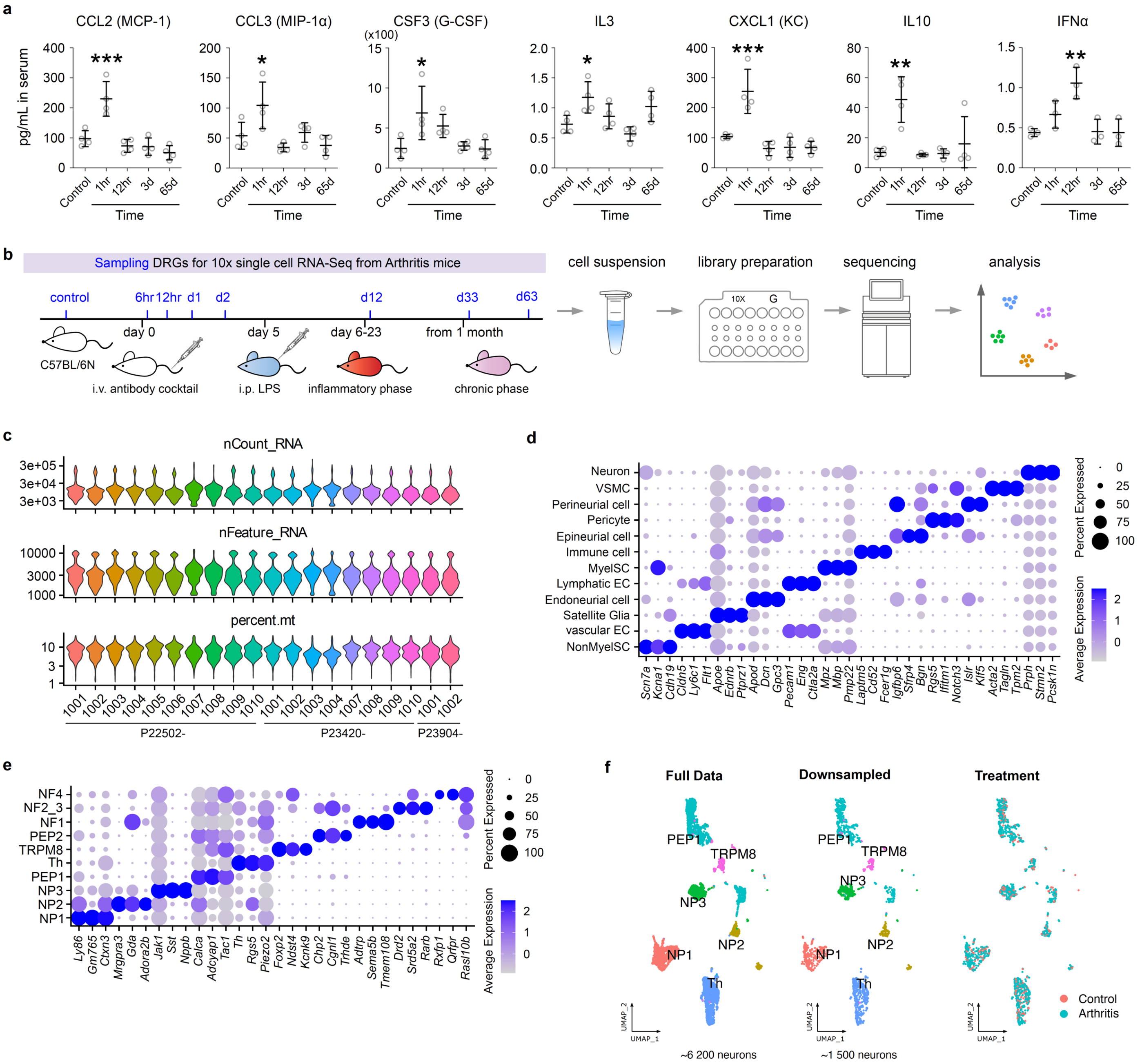


**Extended Data Fig. 3 Single-cell RNA sequencing of DRG neurons in cartilage autoantibody-induced arthritis mice.** (**a**) Cytokine expression in sera collected at different times (hr, hours; d, days) after autoantibody injection and from control mice (n = 4 per group, one-way ANOVA followed by Dunnett multiple comparisons test, **p*<0.05, ***p*<0.01, ****p*<0.001). (**b**) Illustration of sampling and workflow for scRNA-seq of DRG cells from control and autoantibody injected mice. (**c**) Sample metrics violin plots after initial quality control for scRNA-seq data. (**d**) Dot plot of top 3 marker genes for each major cell type of sequenced cells in DRGs. (**e**) Dot plot of top 3 marker genes for each DRG neuronal cell type. (**f**) UMAP of DRG neuronal clusters and of randomly sampled (25%) dataset used for co-regulated gene module analysis showing intermingled distribution of cells from control and arthritis samples.

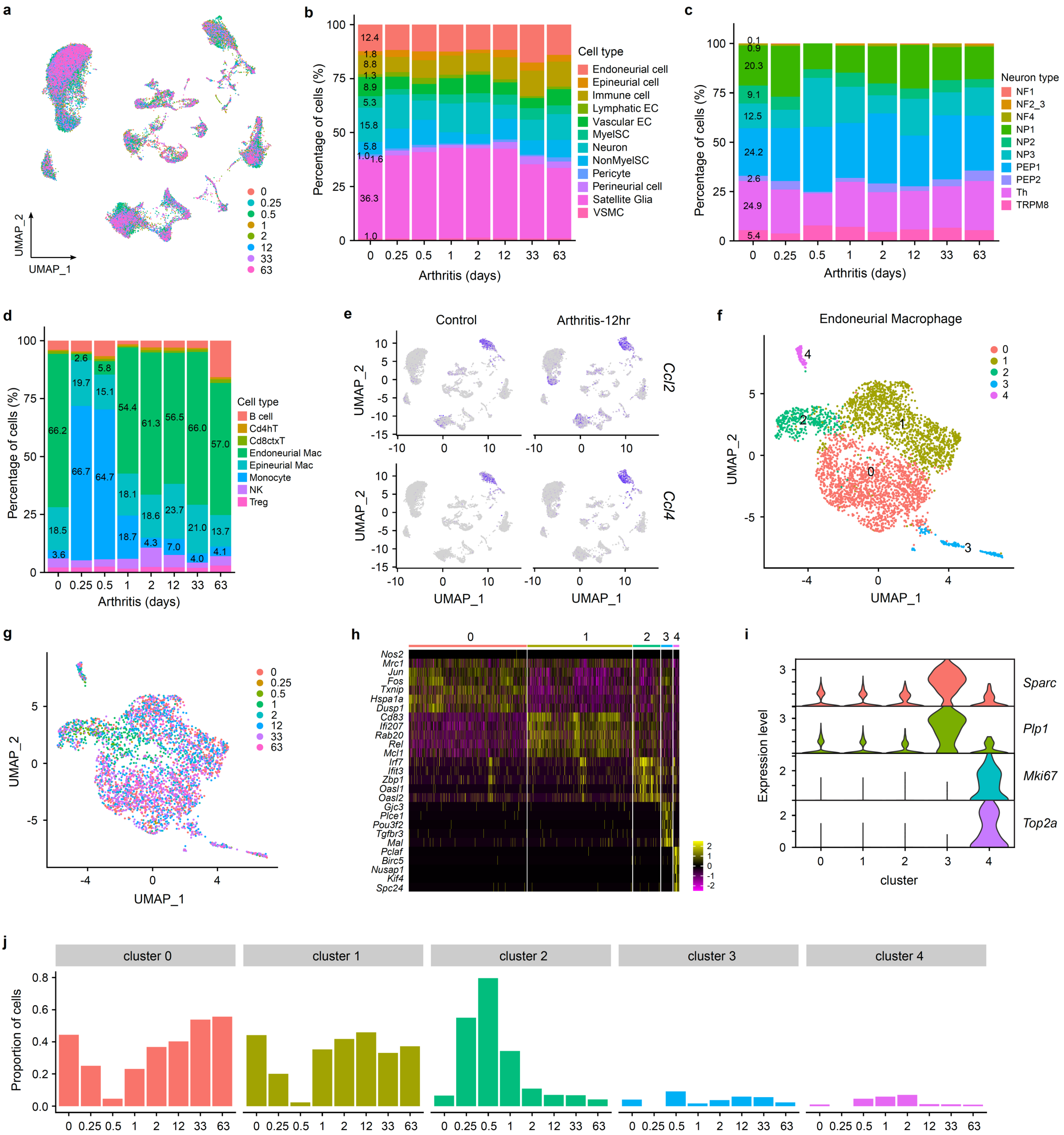

**Extended Data Fig. 4 DRG neuronal and immune cells of autoantibody injected mice identified by sc-RNA seq.** (**a**) UMAP of the distribution of cell clusters (down sampled 5 000 cells from 86 052 cells) from autoantibody injected (0.25, 0.5, 1, 2, 12, 33, 63 days) and control DRG samples. (**b**) Stacked bar plot of the cell type composition (percentage) at different timepoints of after autoantibody injection. (**c**) Stacked bar plot of the neuron type composition (percentage) at different timepoints after autoantibody injection. (**d**) Stacked bar plot of the composition of immune cells (percentage) at different timepoints after autoantibody injection. (**e**) UMAP feature plots for *Ccl2* and *Ccl4* expression in DRGs (12h) from control and autoantibody injected mice show local induced expression in immune cells, non-myelinating Schwann cells and endoneurial macrophages (9 500 cells from each sample). (**f**) UMAP of the distribution of sub-clustering for endoneurial macrophage. (**g**) UMAP of subtypes of endoneurial macrophages showing intermingled distribution of different macrophages from DRGs of control and autoantibody injected mice. (**h**) Heatmap plot shows the top marker genes for each subtype of endoneurial macrophages. (**i**) Stacked violin plot shows marker genes selectively expressed in clusters 3 and 4. (**j**) Dynamic composition of sequenced endoneurial macrophage subtypes in the DRG at different timepoints after autoantibody injection.

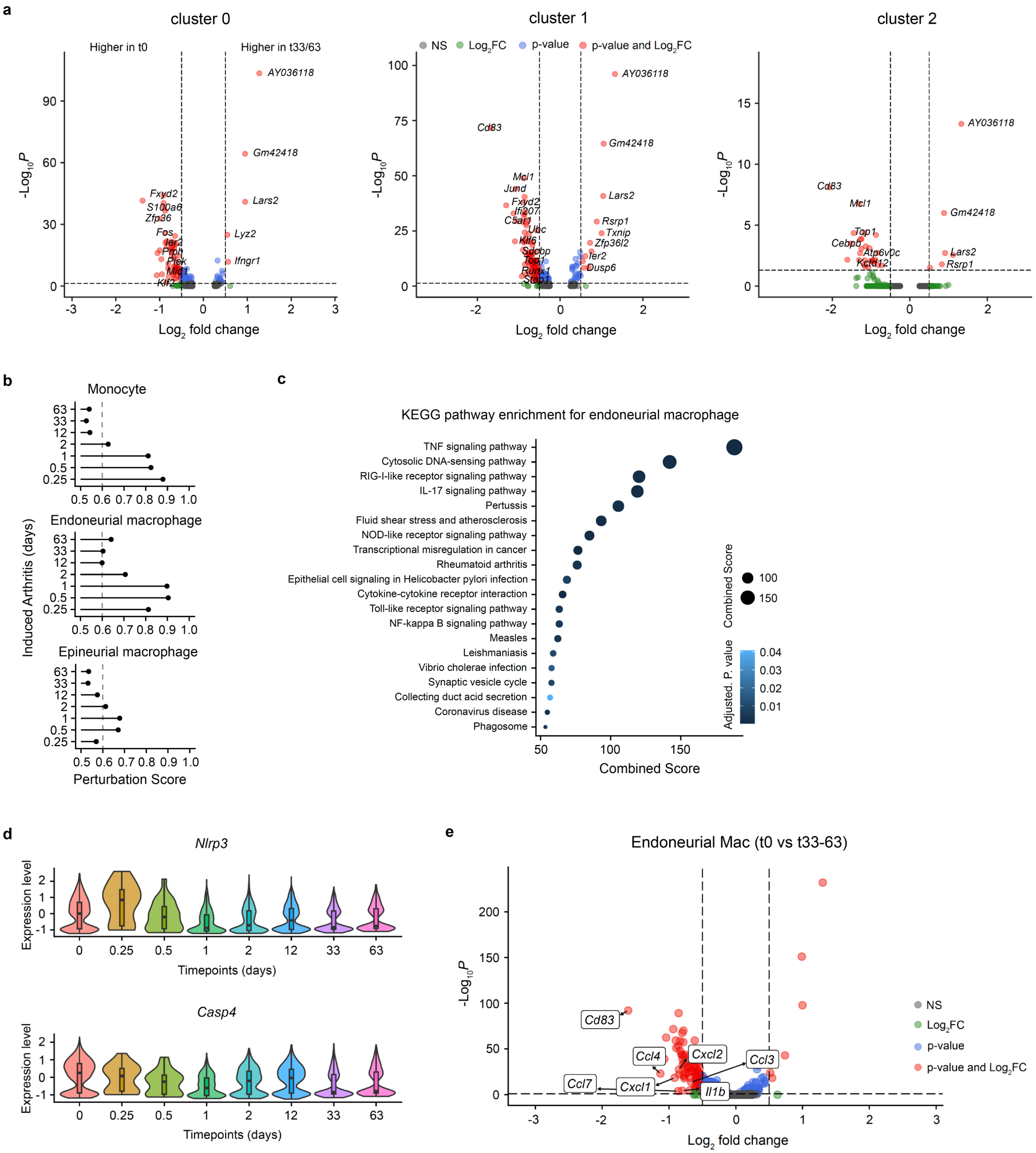


**Extended Data Fig. 5 Differential gene expression analysis for endoneurial macrophages in DRGs of autoantibody injected mice.** (**a**) Volcano plots show up-regulated and down-regulated genes from different subclusters of endoneurial macrophages at the post-inflammatory phase of arthritis (mixed d33 and d63 vs d0). (**b**) Transcriptional perturbation score for non-neuronal DRG cell types (monocytes, endoneurial and epineurial macrophages) at different timepoints after injection of cartilage autoantibodies. (**c**) KEGG pathways of differentially expressed genes in endoneurial macrophages at the post-inflammatory phase of arthritis. (**d**) Violin plots of *Nlrp3* and *Casp4* gene expression in endoneural macrophages of the DRG at different timepoints after administration of autoantibodies to mice. (**e**) Volcano plot of differentially regulated genes from endoneurial macrophages at late (post-inflammatory) phase of arthritis with down-regulated cytokine genes highlighted.

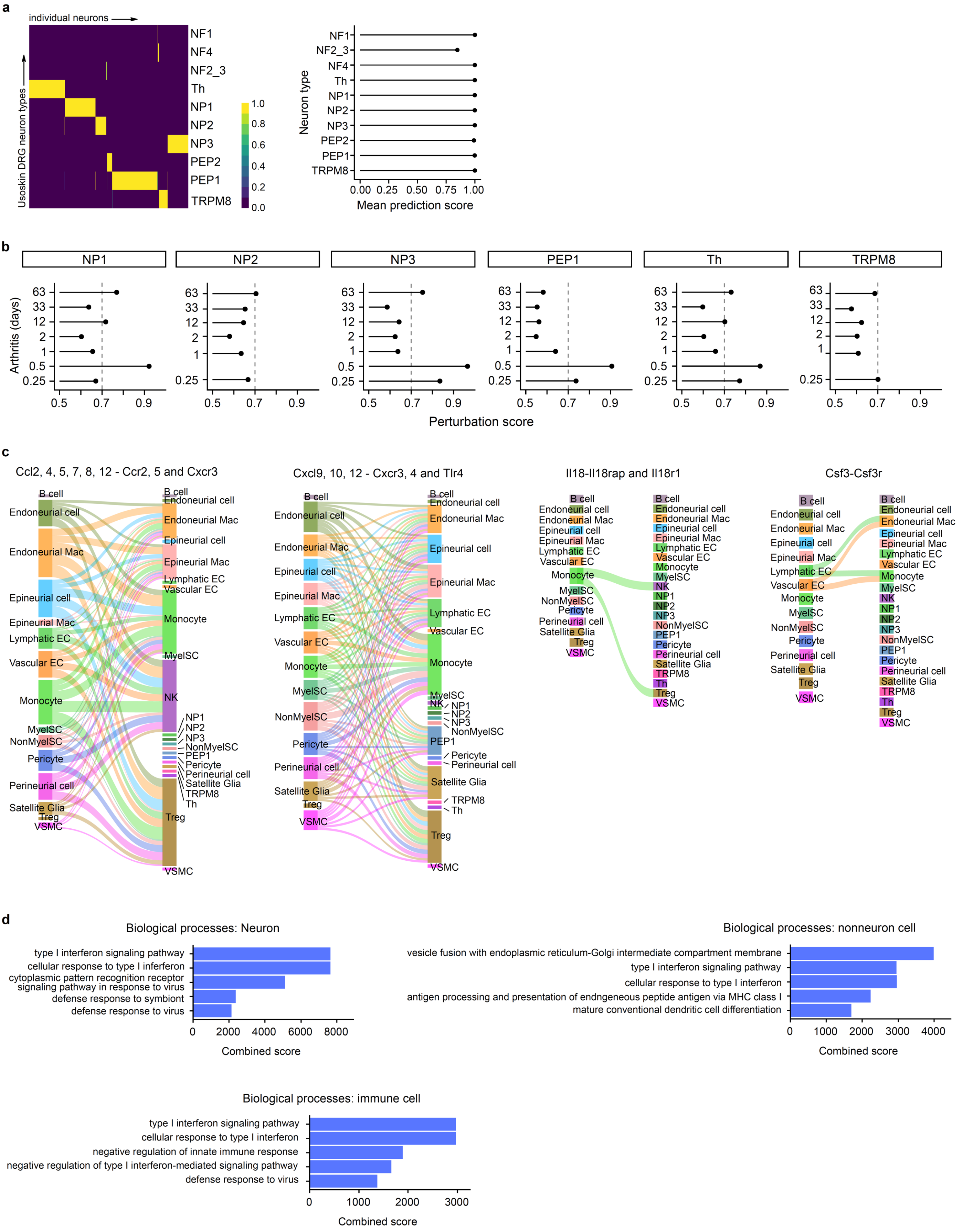


**Extended Data Fig. 6 Transcriptional perturbation analysis for neuronal types and ligand-receptor interaction among DRG cell types.** (**a**) Heatmap (left) representing the predicted similarity score of individual neurons to the neuronal types according to Usoskin's annotation through machine learning classifier. Mean prediction score (right) for different neuronal cell types. (**b**) Perturbation score for DRG neuronal subtypes at different timepoints after injection of autoantibodies in mice. (**c**) Ligand-receptor interactions induced at 12h after autoantibody injection in mice. (**d**) Representative biological processes of differentially expressed genes in neuron, nonneuron and immune cell types.

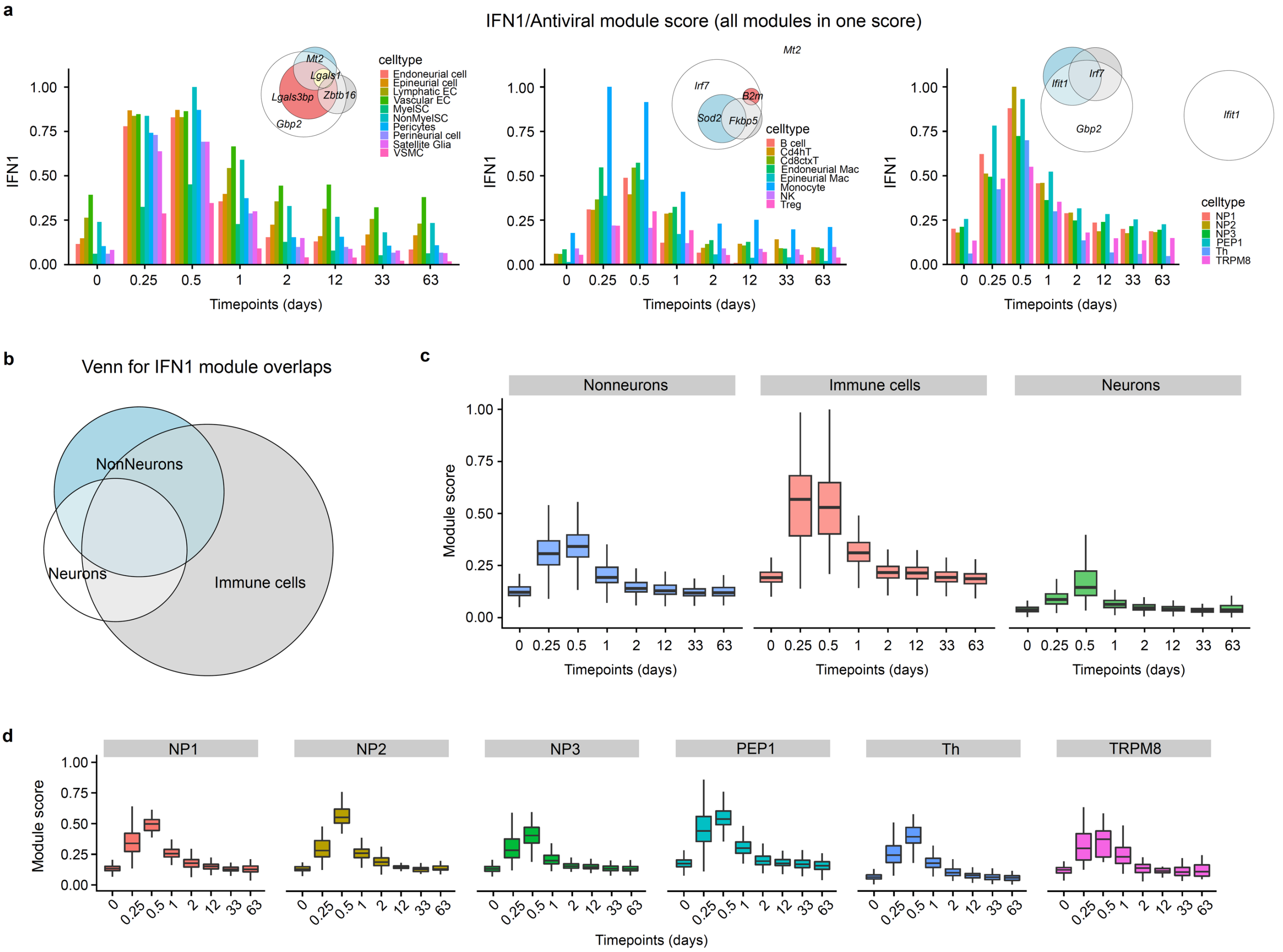


**Extended Data Fig. 7 IFN1/antiviral module gene set enriched by GSEA and co-regulated gene set analysis for DRG cell types in mice with cartilage autoantibodies.** (**a**) IFN1/antiviral co-regulated genes module core for nonneuronal, immune and neuronal cell types at different timepoints after administration of autoantibodies. (**b**) Venn diagram representing the overlapping of IFN1 module genes for nonneurons, neurons and immune cells. (**c**) Boxplots of gene module scores for nonneuronal cells, immune cells and neurons by pseudobulk analysis at different timepoints after autoantibodies were delivered to the mice. (**d**) Boxplots of gene module scores for individual neuronal clusters at different timepoints after autoantibodies were delivered to the mice.

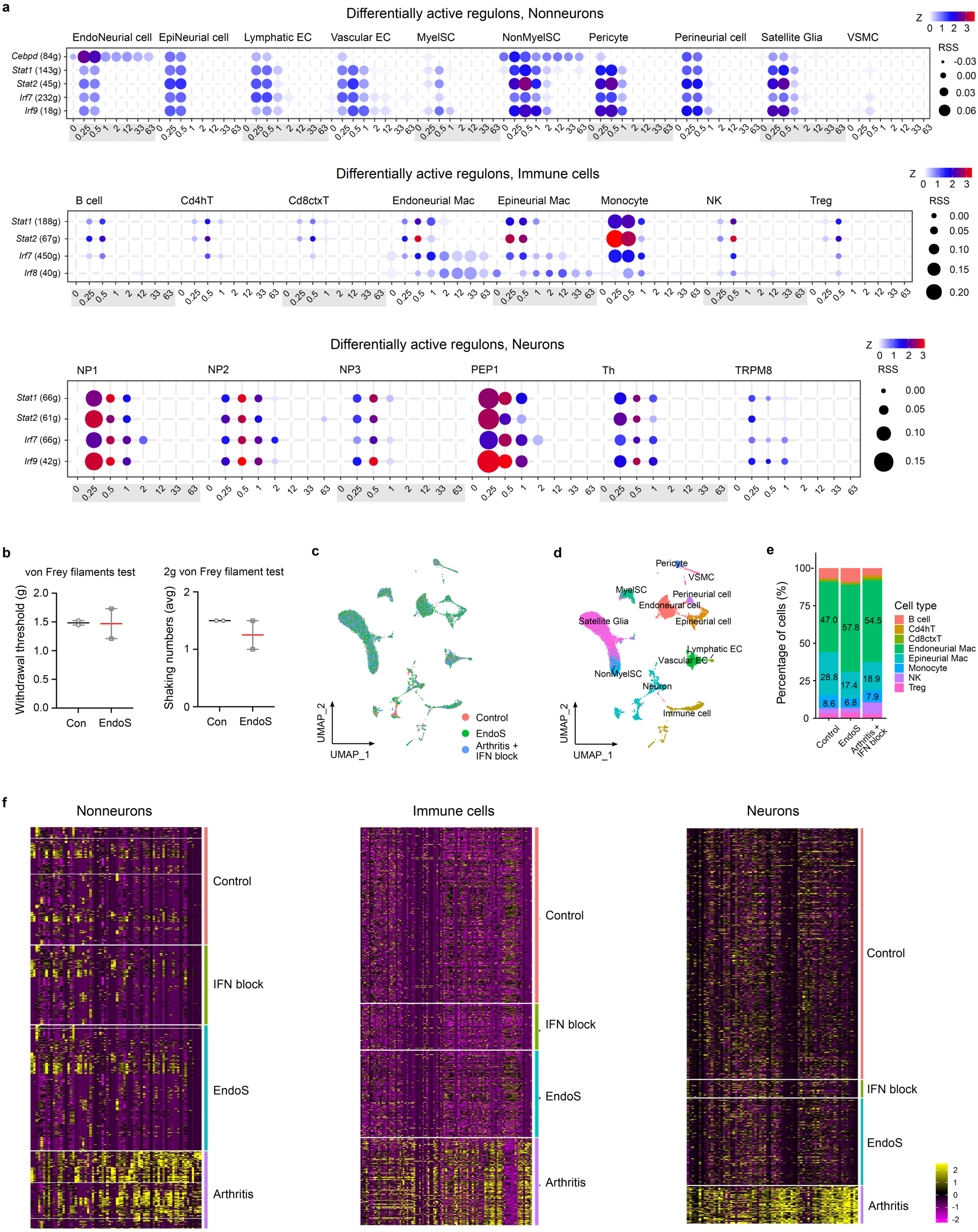


**Extended Data Fig. 8 SCENIC analyses for gene regulatory networks in DRGs and blocking experiment.** (**a**) Dot plots representing enriched IFN-related gene regulons in nonneuronal cells, immune cells and neurons at different timepoints after administration of autoantibodies. (**b**) Mechanical allodynia and hyperalgesia assayed in control and mice injected with EndoS treated autoantibodies (n = 2 per group). Pain-like behavioral tests of von Frey filaments (threshold) and 2g von Frey filament (shaking numbers) show that EndoS treated cartilage autoantibody (EndoS) failed to initiate pain (12h). (**c**) UMAP of sequenced DRG cells showing intermingled distribution of cells from control, EndoS treated autoantibodies and autoantibodies (Arthritis) combined with in vivo IFN block (IFNAR1 antibody). (**d**) UMAP of annotated DRG cell types from sequenced DRG samples in (**c**). (**e**) Stacked bar plot of the composition of immune cells (percentage) from different DRG samples. (**f**) Heatmaps for IFN1 module genes for nonneuronal cell, immune cells and neurons at 0.5 days from control mice, mice pretreated with IFNAR1 antibody and injected with cartilage auto-antibody (IFN block), mice injected with EndoS-treated cartilage autoantibody (EndoS) and mice injected with cartilage autoantibody (Arthritis).

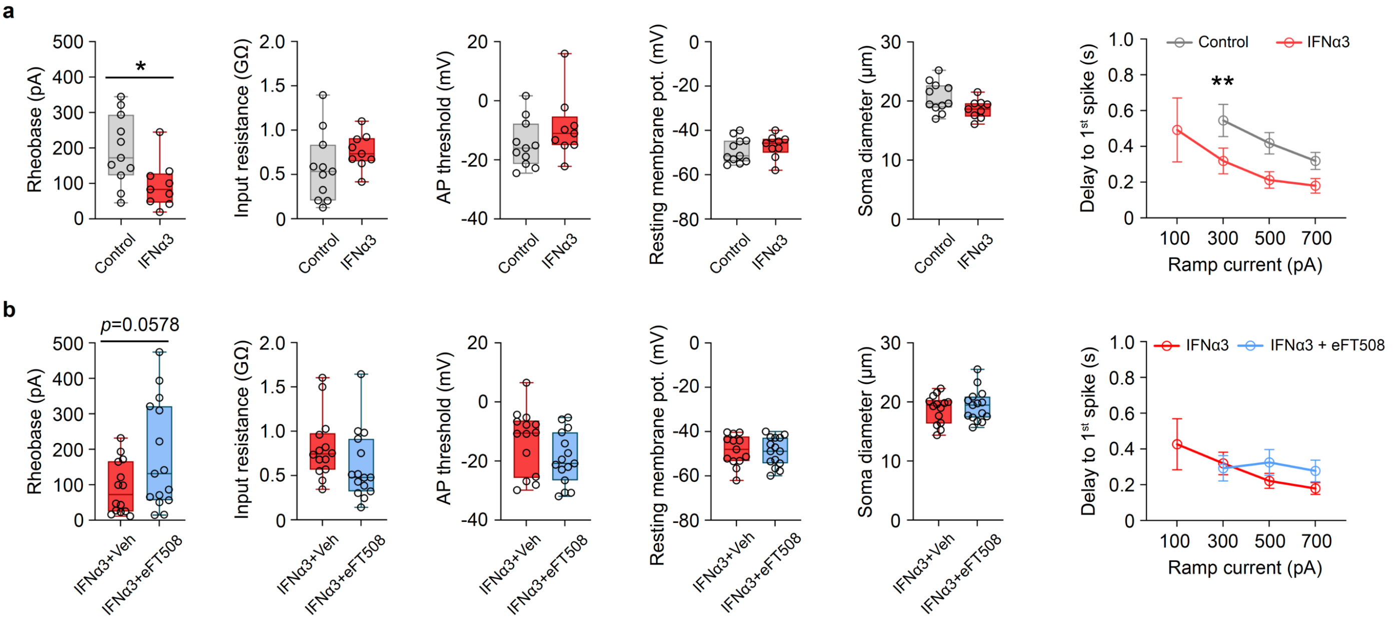


**Extended Data Fig. 9 Patch-clamp recordings of cultured DRG neurons with IFN.** (**a**) Current clamp recording parameters of rheobase, input resistance, action potential threshold, resting membrane potential, soma diameter and the delay to 1st spike at different ramp currents for cultured DRG neurons with no treatment (Control, n = 11) and IFNα3 stimulation (IFNα3, 300 U/ mL, n = 9). (**b**) The same parameter analyses as in (**a**) from cultured DRG neurons treated with IFNα3 + vehicle (DMSO, n = 14) or IFNα3 + eFT508 (MNK1/2 inhibitor, n = 15). Delay to 1st spike data was analyzed with two-way ANOVA followed by Šídák's multiple comparisons test, the rest were analyzed with unpaired *t* test, *$p<0.05$, **$p<0.01$.

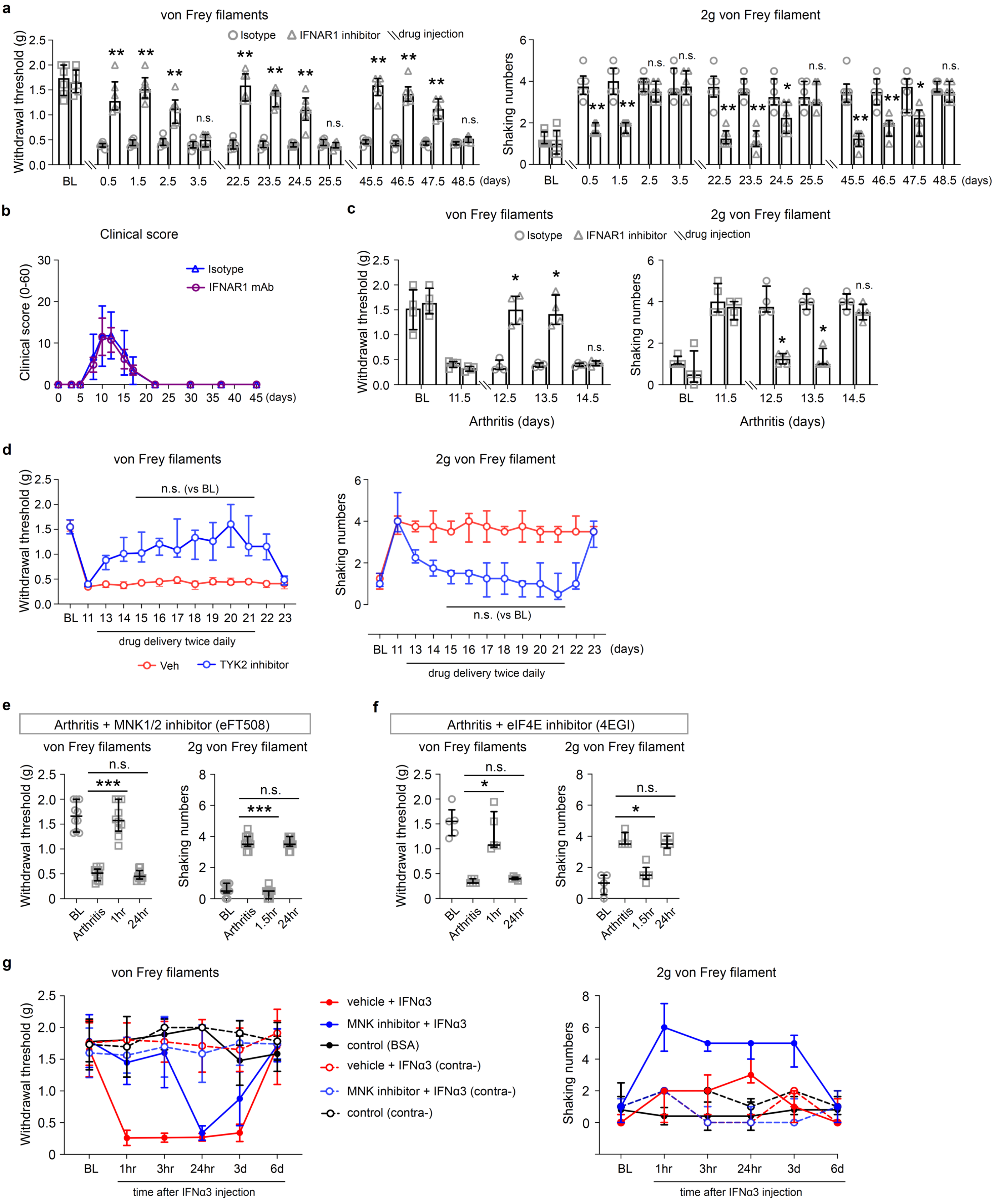


**Extended Data Fig. 10 Sustained interferon signaling drives pain in arthritis.** (**a**) Systemic administration of IFNAR1 mAb (40 mg/kg, i.p.) one hour prior to cartilage autoantibody injection prevented mechanical allodynia (von Frey filaments) and mechanical hyperalgesia (2g von Frey pricking pain behavior) and administration on day 22 and day 45 reversed established allodynia and hyperalgesia compared to isotype IgG1 group (n = 6 per group, **p*<0.05, ***p*<0.01). (**b**) Effects of treatment of IFNAR1 mAb (40 mg/kg, i.p.) as compared to isotype IgG1 administered prior to cartilage autoantibodies on joint inflammation represented as clinical score. (**c**) IFNAR1 mAb administration on day 12 reversed mechanical allodynia and hyperalgesia in arthritis mice with peak inflammation (n = 4 per group, **p*<0.05). (**d**) Reversal of arthritis-induced allodynia and hyperalgesia by TYK2 inhibitor treatment. Five oral administrations of TYK2 inhibitor (Deucravacitinib, 15 mg/kg, twice daily, day13-day21, n = 5-6 per group). (**e**) MNK1/2 inhibitor (eFT508, i.p., 1mg/kg) blocked arthritis-induced allodynia and hyperalgesia in the post-inflammatory phase (n = 10, BL, behavioral response prior to administration of cartilage antibodies; Arthritis, mice at the post-inflammatory phase after cartilage autoantibody administration; 1hr, 1 hr after administration of eFT508 to mice with arthritis at the post-inflammatory phase after cartilage autoantibody administration; 24hr, 24 hr after administration of eFT508 to mice with arthritis at the post-inflammatory phase after cartilage autoantibody administration, ***$p<0.001$). (**f**) eIF4E inhibitor (4EGI, i.p., 15 mg/kg) reversed allodynia and hyperalgesia in the post-inflammatory phase of arthritis (n = 6, BL, behavioral response prior to administration of cartilage antibodies; Arthritis, mice at the post-inflammatory phase after cartilage autoantibody administration; 1.5hr, 1.5hr after administration of 4EGI to mice with arthritis at the post-inflammatory phase after cartilage autoantibody administration; 24hr, 24hr after administration of 4EGI to mice with arthritis at the post-inflammatory phase, *$p<0.05$). (**g**) Pretreatment of MNK1/2 inhibitor eFT508 (2 mg/kg, intraplantar injection) prevented IFN$\alpha$3 (300U, intraplantar) induced mechanical allodynia and hyperalgesia (n = 5 per group). Von Frey filaments and 2g von Frey filament tests were analyzed with Kruskal-Wallis test followed by Dunn's multiple comparisons test.

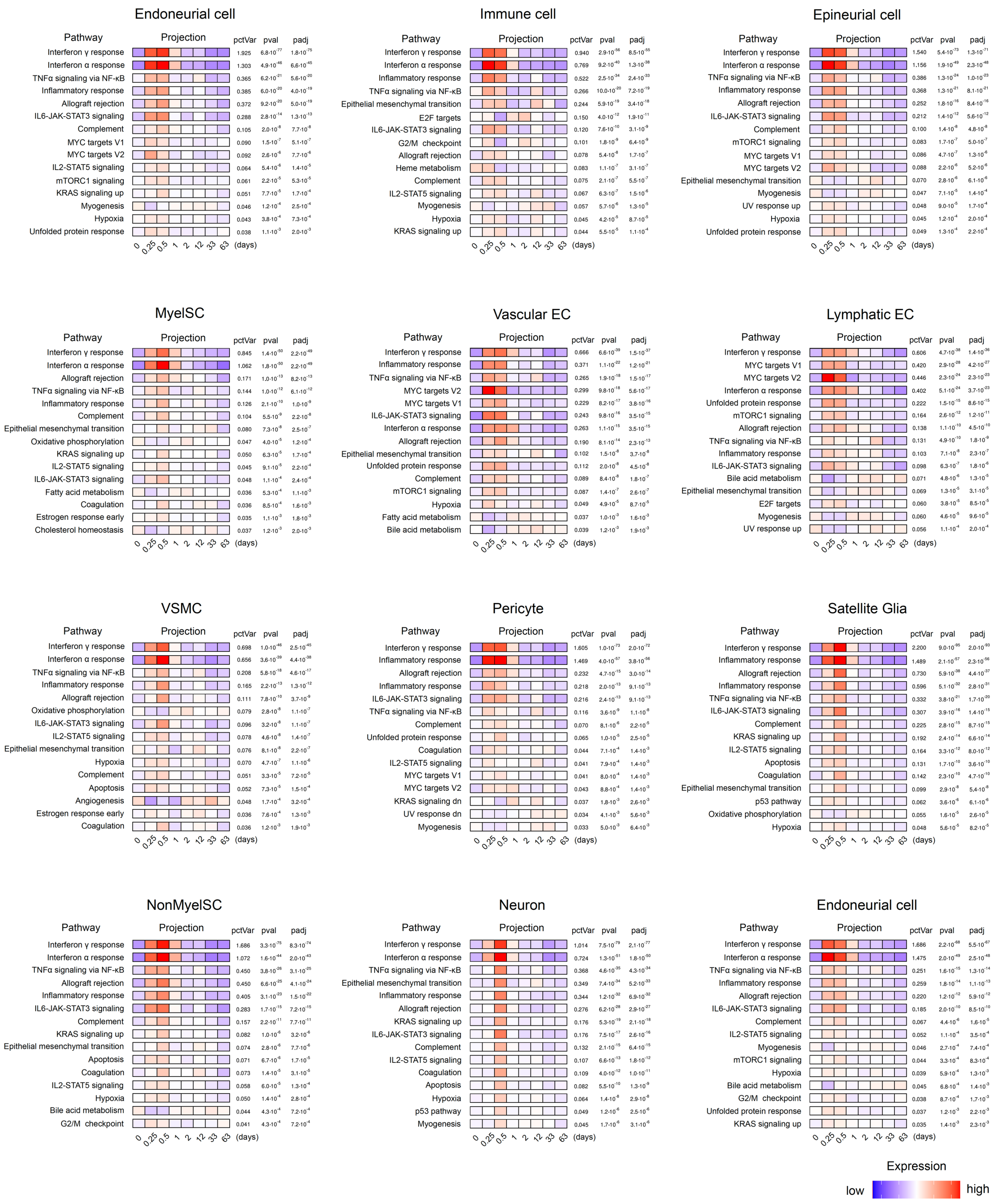


**Supplementary Fig. 1 Gene-set enrichment analysis (GSEA) of DRG cell types.** GSEA analysis for hallmark genes in DRG cell types at different timepoints after injection of cartilage autoantibodies in mice.

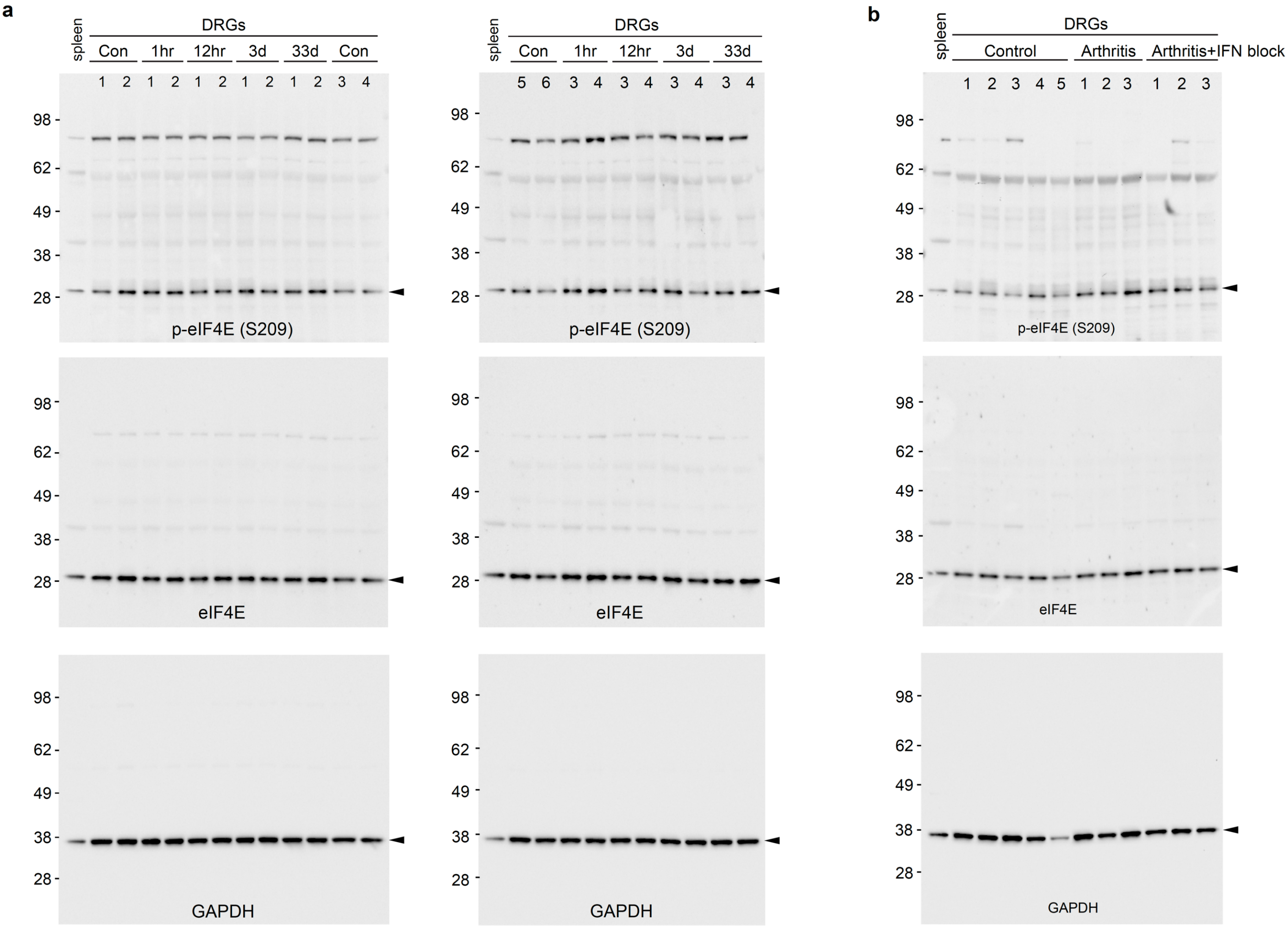


**Supplementary Fig. 2 Western blotting of phospho-eIF4E (Ser209) in DRGs of cartilage autoantibody injected mice.** (**a**) Western blotting of phospho-eIF4E (Ser209), eIF4E and GAPDH in DRG lysates from control and cartilage autoantibody injected mice at different timepoints. (**b**) Western blotting of phospho-eIF4E (Ser209), eIF4E and GAPDH in DRG lysates from control, arthritis mice (d33) and arthritis mice treated with IFNAR1 antibody (IFN block). Arrowhead indicates the specific band. Spleen sample was added as positive control. DRG samples from different mice are numbered at the top of the membranes.

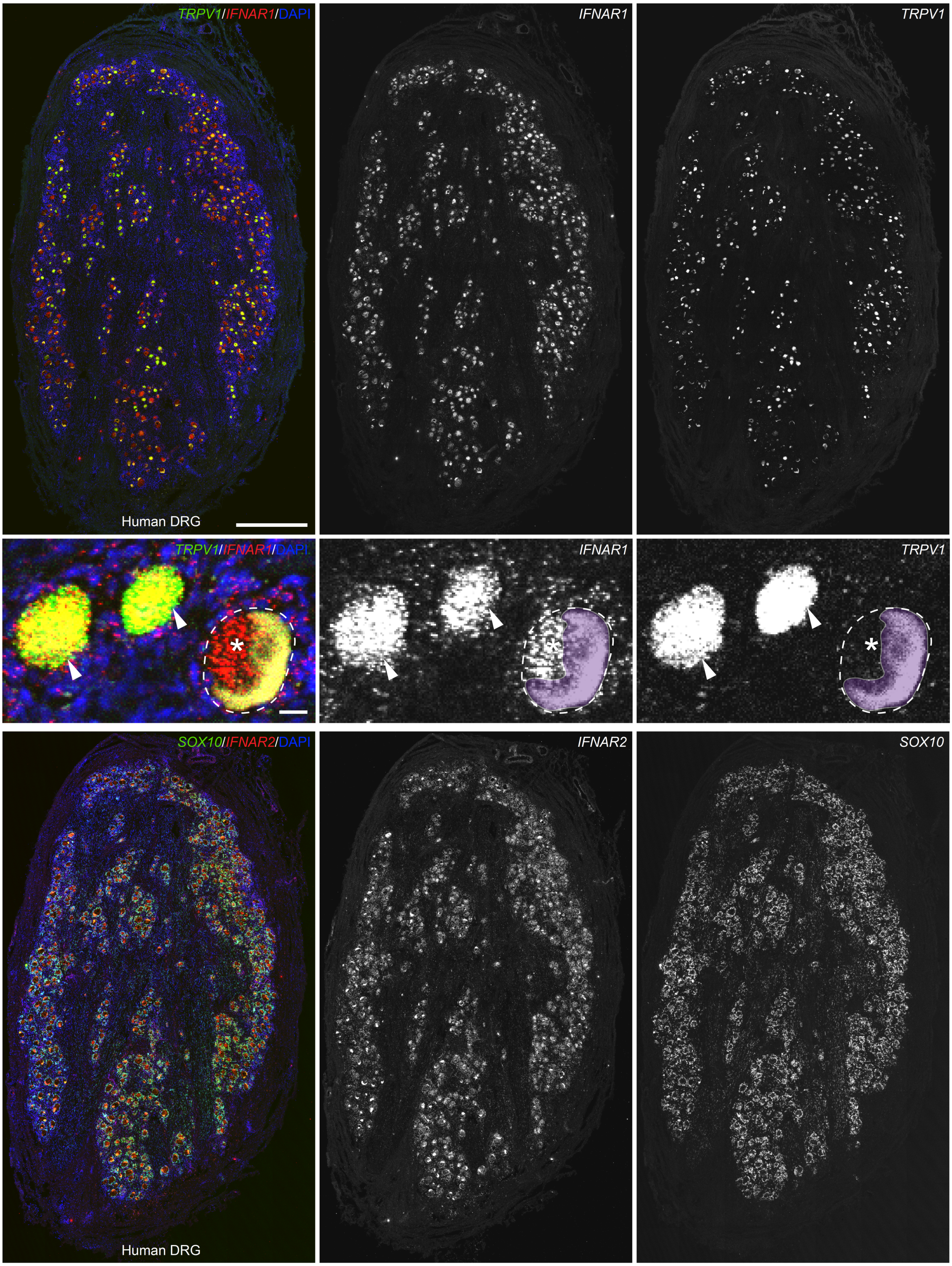


**Supplementary Fig. 3 Expression of IFNAR1 and IFNAR2 mRNAs in human DRGs detected by RNAscope.** Top: Representative overview image of co-expression of *TRPV1* and *IFNAR1* mRNAs in human DRG counter stained with DAPI. Middle: High magnification image of co-localization between *TRPV1* and *IFNAR1* mRNAs in DRG neurons as indicated by arrowheads. The asterisk indicates autofluorescence caused by accumulated lipofuscin. Bottom: Representative overview image of *SOX10* and *IFNAR2* mRNA expression in human DRG counter stained with DAPI. Scale bars: 1 mm for overview images on top and bottom panels and 20 μm for middle panel.
