## Supplementary Tables and Videos for "Persistent interferon signaling that causes sensory neuron plasticity and pain in arthritis": Supplementary Table_8_Information_of_human_DRG_tissue.docx

| Sample | Sex | Age | Ethnicity | BMI | Pain |
| --- | --- | --- | --- | --- | --- |
| Con1 | Female | 43 | Caucasian | 26,2 | no |
| Con2 | Female | 32 | Caucasian | 23,9 | no |
| Con3 | Female | 31 | Caucasian | 28,3 | no |
| Con4 | Female | 38 | Caucasian | 26,6 | no |
| Con5 | Female | 51 | Caucasian | 20,3 | no |
| Con6 | Female | 45 | African Ameican | 20,3 | no |
| Con7 | Female | 41 | African Ameican | 28 | no |
| Con8 | Female | 44 | Caucasian | 30,9 | no |
| Con9 | Female | 32 | Caucasian | 25,8 | no |
| Con10 | Female | 46 | Caucasian | 31,7 | no |
| Con11 | Female | 47 | Hispanic | 29 | no |
| RA1 | Female | 62 | Caucasian | 29,4 | Chronic pain |
| RA2 | Female | 56 | African Ameican | 22,5 | Chronic pain |
| RA3 | Female | 46 | Caucasian | 21 | Chronic pain |
| RA4 | Female | 61 | Caucasian | 32,3 | no |
| RA5 | Female | 34 | African Ameican | 41,8 | no |
| RA6 | Female | 37 | African Ameican | 30,4 | no |
| RA7 | Female | 59 | Caucasian | 20,8 | no |
| RA8 | Female | 37 | Caucasian | 15,8 | no |

Supplemental Table S8. Information of donors for DRG tissues. Lumbar DRGs (L3-L5) from rheumatoid arthritis patients (3 RA with pain; and 5 RA without pain) and healthy controls (matched, n=11).
